# Role of the reaction-structure coupling in temperature compensation of the KaiABC circadian rhythm

**DOI:** 10.1101/2021.10.11.464015

**Authors:** Masaki Sasai

## Abstract

When the mixture solution of cyanobacterial proteins, KaiA, KaiB, and KaiC, is incubated with ATP in vitro, the phosphorylation level of KaiC shows stable oscillations with the temperature-compensated circadian period. Elucidating this temperature compensation is essential for understanding the KaiABC circadian clock, but its mechanism has remained a mystery. We analyzed the KaiABC temperature compensation by developing a theoretical model describing the feedback relations among reactions and structural transitions in the KaiC molecule. The model showed that the reduced structural cooperativity should weaken the negative feedback coupling among reactions and structural transitions, which enlarges the oscillation amplitude and period, explaining the observed significant period extension upon single amino-acid residue substitution. We propose that an increase in thermal fluctuations similarly attenuates the reaction-structure feedback, explaining the temperature compensation in the KaiABC clock. The model explained the experimentally observed responses of the oscillation phase to the temperature shift or the ADP-concentration change and suggested that the ATPase reactions in the CI domain of KaiC affect the period depending on how the reaction rates are modulated. The KaiABC clock provides a unique opportunity to analyze how the reaction-structure coupling regulates the system-level synchronized oscillations of molecules.

**Author summary:** The reconstituted KaiABC circadian clock provides a unique opportunity to analyze how the effects of chemical and structural features of individual molecules determine the system-level oscillations of many molecules. By modeling the coupling of chemical reactions and structural transitions in the KaiC molecule, we showed that reducing the coupling strength enlarges the oscillation amplitude and period, explaining the observed striking change of the period length upon single-residue substitution in KaiC. We propose that thermal fluctuations attenuate the reaction-structure coupling similarly to the residue substitution, explaining the stable temperature compensation observed in the KaiABC clock. The combined experimental and theoretical analyses should open a way to develop techniques to design the system-level molecular oscillations, further providing a basis for understanding circadian clocks in vivo.

## Introduction

The mixture solution of cyanobacterial proteins, KaiA, KaiB, and KaiC, shows the robust structural and chemical oscillations with the period of approximately 24 h when the solution is incubated with ATP in vitro [1, 2, 3]. This period is insensitive to the temperature change, showing the feature of temperature compensation [1]. An important question is whether this feature has a common molecular mechanism or the same mathematical principle as temperature compensation in generic transcription-translation oscillations (TTO). Circadian clocks in many organisms are driven by the time-delayed negative feedback in the TTO [4, 5], whose oscillation period is temperature compensated [6, 7]; the ratio of the period in 10 °C difference is 0.9 ≲ *Q*_10_ ≲ 1.1, which is much closer to 1 than the value ≳ 1.5 expected from the temperature dependence of normal biochemical reactions. This temperature compensation has been studied with various theoretical models [8, 9, 10, 11, 12, 13, 14, 15, 16, 17, 18, 19], but its mechanism remains a challenging problem.

There have been at least three views or approaches to temperature compensation; (i) the balance between opposing reactions, (ii) the correlation between oscillation period and amplitude, and (iii) the role of a critical reaction step. One view is the balance between reactions opposingly working to shorten or lengthen the period upon temperature change [7, 9]. Such balancing was mathematically formulated [20] and studied with different models [10, 11, 12, 13, 14, 15]. In searching for the balancing reactions, various mechanisms were examined, including the balance between negative and positive feedback regulations [9] and the balance between ways of resetting bifurcations [12]. In particular, Lakin-Thomas et al. emphasized that the period length should correlate with the amplitude of oscillations [8], suggesting the possible use of the correlation as a clue to find the reactions responsible for the balance [17]. An alternative approach is to search for the critical reaction step or the molecule that determines temperature compensation. In mammals, phosphorylation of period 2 (PER2) regulated by casein kinase I*ε*/*δ* (CKI*ε*/*δ*) is temperature insensitive [21], and this insensitivity was attributed to the reaction mechanism of the CKI*ε*/*δ* molecule [22].

Therefore, it is meaningful to analyze the temperature compensation of the KaiABC post-translational oscillations from the three views discussed for TTO. Previously, two hypotheses on the KaiABC temperature compensation were proposed based on the view (iii) of the critical reaction step and the view (i) on the balance between competing reactions. A critical reaction step is ATP hydrolysis in KaiC. KaiC hexamer is a slow ATPase, and variation of the ATPase activity among KaiC mutants is correlated with the variation of the oscillation frequency of those mutants [23, 24]. Because the ATPase activity of KaiC is temperature insensitive [23], this correlation suggested that ATPase activity determines temperature compensation in the KaiABC oscillations. In order to clarify whether such causality exists behind the observed correlation, further experimental and theoretical investigations are necessary. Another hypothesis was based on the assumption of balance among the competing binding reactions of KaiA to KaiC at different phosphorylation levels [25]. With this hypothesis, the population of KaiC at a highly phosphorylated state dominates in low temperature, which increases the free unbound KaiA molecules, and increases the overall binding rate of KaiA to compensate for the decrease of reaction rates at the low temperature. However, the accumulation of KaiC at a highly phosphorylated state in low temperature was not observed experimentally [1]; therefore, exploration for other hypotheses and comparison among them are necessary.

Here, we propose a hypothesis based on the view (ii) of the correlation between oscillation period and amplitude. A recent experimental report revealed that substituting an amino-acid residue near the CI and CII domains interface in KaiC induces a striking change in the period from 15 h to 158 h with a suggested correlation between period and amplitude [26]. The modified period length in the mutant was anti-correlated with the volume of the residue after the substitution [26], indicating that the structural coupling between the CI and CII domains is crucial to determining the period; the period was enlarged when the structural coupling between domains was weaker with the smaller volume of residue at the domain interface. The recent X-ray analysis showed that KaiC undergoes structural transitions depending on the state of two phosphorylation sites in the CII and the nucleotide-binding state in the CI [27]. This observation indicated that the structural transitions of KaiC take place through the allosteric communication between CI and CII domains depending on the phosphorylation reactions in the CII and the ATPase reactions in the CI. On the other hand, the rates of those reactions should depend on the structure. Therefore, it is plausible to assume the feedback relations among reactions and structural transitions. Substituting smaller volume residue at the CI-CII interface should reduce the transition cooperativity and weaken the feedback coupling. The weakening of the negative feedback lengthens the period in general nonlinear oscillators as found in the TTO model [9]; therefore, the observed change in the mutants can be explained if the coupling among reactions and structural transitions of KaiC constitutes the negative feedback relation. Modifying strength of the negative feedback leads to the correlated change in amplitude and period in general nonlinear oscillators [9]. Ito-Miwa et al. compared five examples of the CI-CII interface mutations, showing two of them have smaller amplitude with shorter period [26], consistently with the negative-feedback hypothesis. However, the amplitude change of the rest three mutations was not sufficiently quantified [26], suggesting the need for the further statistical evaluation of the mutant oscillations.

In the present study, we propose that the structural coupling between the CI and CII is weakened through thermal fluctuations, weakening the negative feedback relations. In the higher temperature, the larger thermal fluctuations at the CI-CII interface should obscure the specific atomic interactions at the interface, producing a similar effect to the substitution to the smaller volume residue. This weakening of interactions at the interface reduces the negative feedback strength, enhancing the oscillation amplitude, lengthening the period, and compensating for the thermal acceleration of reactions in the higher temperature. We analyze this hypothesis with a model of the KaiABC oscillator and discuss possible tests of the model prediction. We also analyze the correlation between the ATPase reactions and the oscillation frequency to discuss the role of the ATPase reactions in the Kai system temperature compensation.

## Model

### Problems at two levels; the molecular and ensemble levels

In modeling the KaiABC oscillator, we need to analyze two mechanisms: how individual KaiC molecules oscillate and how oscillations of many KaiC molecules synchronize to generate the ensemble-level oscillations in solution. Many theoretical works focused on the latter question as the sequential change of phosphorylation level of individual KaiC molecules was assumed in advance; then, the synchronization was explained using various hypotheses [28, 29, 30, 31, 32, 33, 34, 35, 25, 36, 37, 38, 39].

A plausible hypothesis is based on KaiA sequestration [34, 35, 37, 38, 25, 39, 40, 36, 41, 42, 43, 44]; the preferential KaiA binding to particular KaiC states sequestrates KaiA, reducing the KaiA binding rate in the other KaiC states, leading to the accumulation of the population in those states, and producing coherent synchronizaed oscillations. This hypothesis is consistent with the experimental observation that the ensemble oscillations disappear when KaiA is too abundant in the solution [45]. Various states of KaiC were assumed as the KaiA-sequestrating states; some models assumed KaiA is sequestrated into the lowly phosphorylated KaiC in the phosphorylation (P) process [34, 35, 25, 40, 36] or in the dephosphorylation (dP) process [37, 38]. The other models assumed that KaiA is sequestrated by forming the KaiC-KaiB-KaiA complexes that appear during the dP process [39, 41, 42, 43, 44]. The present author showed [44] that the assumption of the KaiA sequestration into the KaiC-KaiB-KaiA complexes quantitatively explains the experimental data on how the oscillations are entrained when two solutions oscillating at different phases are mixed [46].

In the present study, we use the hypotheses of the KaiA sequestration into the KaiC-KaiB-KaiA complexes to explain the synchronization. We also model the mechanism of how oscillations are driven in individual KaiC molecules. In this way, we address the questions extending over the two levels, the individual molecular level and the ensemble level, to analyze how the molecular-level feedback coupling determines the ensemble-level oscillations and temperature compensation.

### Feedback coupling of reactions and structural transitions at the molecular level

At the molecular level, KaiC forms a hexamer [47, 48, 49], which is denoted here by C_6_. The KaiC monomer is composed of the N-terminal (CI) and C-terminal (CII) domains [50], which are assembled to the CI and CII rings in C_6_ [49] (Fig. 1). The NMR [51, 52], small-angle X-ray diffraction [53], biochemical analyses [54], and X-ray crystallography [27] showed the cooperative structural transitions of KaiC hexamer between two, the structure in the P phase and the structure in the dP phase. We use the order parameter 0 ≤ *X* ≤ 1 to describe the transitions between two typical conformations and thermal fluctuations around each conformation. We write *X*(*k,t*) ≈ 1 when the *k*th KaiC hexamer at time *t* takes the structure in the P phase, and *X*(*k,t*) ≈ 0 when it takes the structure in the dP phase

**Figure 1:**
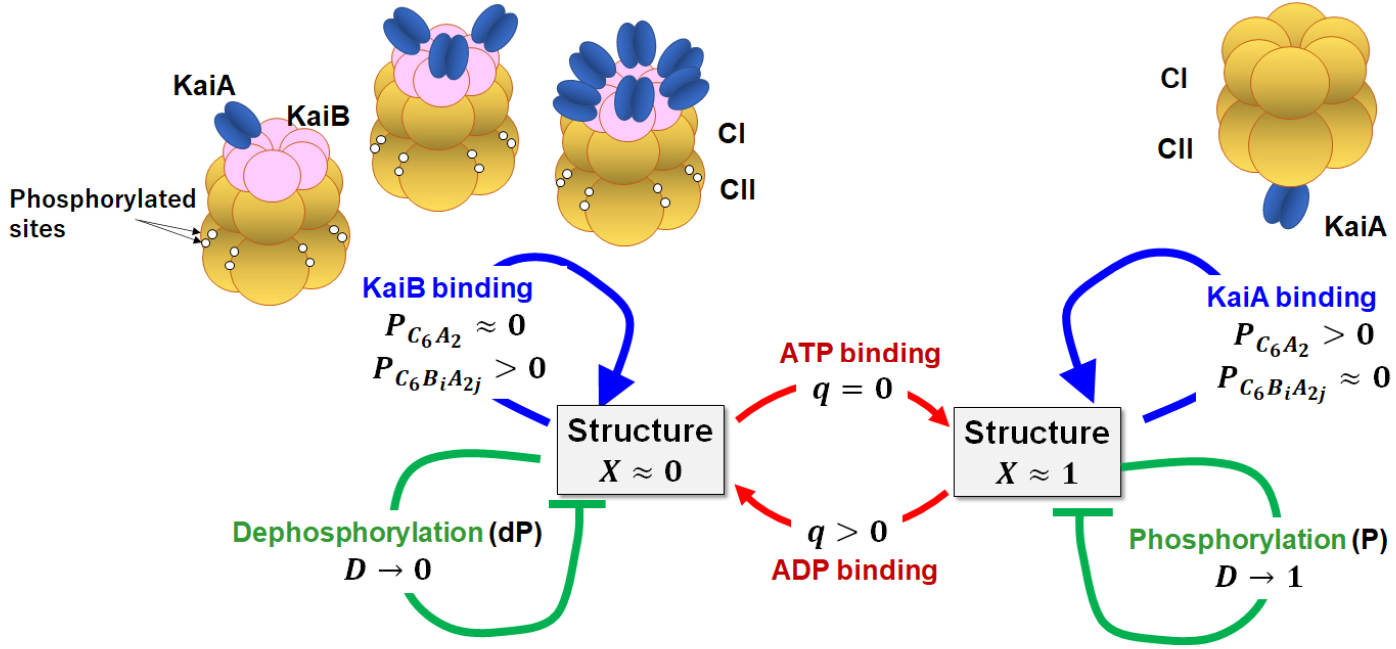
Feedback coupling among reactions and structural transitions in KaiC hexamer. The model is based on the fundamental experimental observations: KaiC forms a hexamer composed of the CI and CII rings. The CII has twelve sites to be phosphorylated. The KaiC hexamer undergoes allosteric transitions between two structures; the structure (*X* ≈ 1) in the phosphorylation (P) phase and the structure (*X* ≈ 0) in the dephosphorylation (dP) phase. A KaiA dimer can bind on the CII of the *X* ≈ 1 KaiC with the probability *P*_C_6_A_2__, which promotes the P process to increase the phosphorylation level *D.* KaiB monomers can bind on the CI of the *X* ≈ 0 KaiC, and a KaiA dimer can further bind on each KaiB monomer forming KaiC-KaiB-KaiA complexes with the probability *P*_C_6_B_*i*_A_2*j*__ with *j* ≤ *i* ≤ 6. KaiA unbinds from the CII of the *X* ≈ 0 KaiC, promoting the dP process to decrease *D*. Based on these observations, the model describes the feedback coupling in the KaiC hexamer by assuming (**i**) the KaiA binding on the CII stabilizes the *X* ≈ 1 state, and (**ii**) the KaiB binding on the CI stabilizes the *X* ≈ 0 structure. The assumptions (**i**) and (**ii**) account for the positive feedback to stabilize the *X* ≈ 1 and *X* ≈ 0 states. The model also assumes that (**iii**) the gradual rise of D destabilizes the *X* ≈ 1 structure and (**iv**) the gradual fall of *D* destabilizes the *X* ≈ 0 structure. The assumptions **(iii**) and (**iv**) support the time-delayed negative feedback to drive the transitions between the *X* ≈ 1 and *X* ≈ 0 states. The model further assumes (**v**) the stochastic ATP hydrolysis in the CI (*q* → 1) destabilizes the *X* ≈ 1 state, and (**vi**) the stochastic ADP release from the CI and the subsequent ATP binding (*q* → 0) destabilize the *X* ≈ 0 state. (**v**) and (**vi**) trigger the transitions between the *X* ≈ 1 and *X* ≈ 0 states. The assumptions (**i**) through (**vi**) generate the oscillations of individual KaiC hexamers. We hypothesized (**vii**) oscillations of multiple KaiC hexamers are coupled through the KaiA sequestration into the KaiC-KaiB-KaiA complexes, giving rise to the ensemble-level oscillations.

In the present model, we consider that the structural state is a hub of multifold feedback relations among reactions and structural transitions (Fig. 1). We describe individual KaiC molecules with the structural parameter *X* and coarse-grained variables representing three types of reactions; (1) the binding/unbinding reactions of KaiA and KaiB to/from KaiC, (2) the P/dP reactions in the CII, and (3) the ATPase reactions in the CI. We consider that these three types of reactions directly or indirectly depend on X, and these reactions affect *X*, constituting the multifold feedback relations.

#### Binding/unbinding reactions of KaiA and KaiB

KaiA and KaiB bind/unbind to/from KaiC in a coordinated way [55, 56, 57, 58, 59, 60]. The CII ring of a KaiC hexamer can bind a KaiA dimer during the P process to form C_6_A_2_ [61, 36]. The cryo-electron microscopy and mass-spectrometry showed that each CI domain can bind a KaiB monomer to form KaiC-KaiB complexes, further binding KaiA dimers to form KaiC-KaiB-KaiA complexes, C_6_B_*i*_A_2*j*_ with *j* ≤ *i* ≤ 6 [60]. The stoichiometry C_6_B_*i*_A_2*j*_ implies the large capacity of KaiC-KaiB-KaiA complexes to absorb KaiA molecules. We consider the probability of the *k*th KaiC hexamer forming C_6_A_2_ at time *t, P*_C_6_A_2__ (*k,t*), and the probability forming C_6_B_*i*_A_2*j*_, *P*_C_6_B_*i*_A_2*j*__(*k,t*).

Binding/unbinding of KaiA to/from the CII has a timescale of seconds [36]. Because this kinetics is much faster than the other reactions in KaiC, we describe *P*_C_6_A_2__ with the quasi-equilibrium approximation, *P*_C_6_A_2__(*k,t*) = *x*_A_(*t*)*g*_C:A_(*k, t*)*P*_C_6_B_0_A_0__ (*k,t*) with *g*_C:A_(*k,t*) = *h_A_*(*k, t*)/*f*_A_(*k, t*), where *x*_A_(*t*) is the concentration of free unbound KaiA dimer at time t. *h*_A_(*k,t*) and *f*_A_(*k,t*) are the binding and unbinding rate constants of a KaiA dimer to and from the CII, respectively. We represent the preferential KaiA binding to the *X* ≈ 1 structure by assuming that *h*_A_(*k,t*) is an increasing function and *f*_A_(*k,t*) is a decreasing function of *X*(*k,t*) (See Methods).

Binding/unbinding of KaiB has a timescale of an hour [39, 36]. We describe the slow temporal variation of *P*_C_6_B_*i*_A_2*j*__(*k,t*) by integrating the kinetic equations (Methods), which are represented with the rate constants of KaiB binding and unbinding, *h*_B_(*k,t*) and *f*_B_(*k,t*), respectively, and the rate constants of KaiA binding and unbinding to and from the KaiB, *h*_AB_ and *f*_AB_, respectively. We represent the tendency of preferential binding of KaiB to the *X* ≈ 0 structure by assuming that *h*_B_ is a decreasing function and *f*_B_ is an increasing function of *X*(*k,t*). Because KaiA does not directly interact with KaiC in this process, we assume *h*_AB_ and *f*_AB_ are independent of *X*. The unusually slow yet specific binding kinetics of the KaiB should be attributed to the fold transitions of KaiB. KaiB switches between ground state (KaiB_gs_) and fold-switched state (KaiB_fs_) by changing its secondary structures [62]. When only KaiB_fs_ has a significant binding affinity to the CI and KaiB_fs_ is the excited state with the activation energy of Δ *E*_gs→fs_ > 0, the small factor exp(-Δ*E*_gs→fs_/(*k*_B_*T*)) explains the small *h*_B_.

#### P/dP reactions

Each CII domain has two sites, Ser431 and Thr432, to be phosphorylated, which amounts to 12 sites in a KaiC hexamer. For simplicity, we do not distinguish Ser431 and Thr432 in the present expression, describing the phosphorylation level of 12 sites with the parameter 0 ≤ *D*(*k,t*) ≤ 1; *D*(*k, t*) = 1 when 12 sites in the CII of the *k*th KaiC hexamer are all phosphorylated, and *D*(*k, t*) = 0 when they are all unphosphorylated. Phosphorylation is promoted when KaiA binds on the CII [55, 39], and dephosporylation proceeds when it unbinds [39]. We represent this tendency by writing

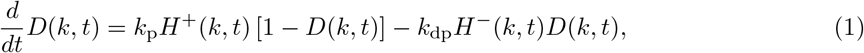

where *H*^+^(*k,t*) = *z*/(1 + *z*) and *H*^-^(*k,t*) = 1/(1 + *z*) represent the effects of binding and unbinding of KaiA with *z* = *P*_C_6_A_2__(*k,t*)/*P*_0_ and a constant *P*_0_. For changing *D* between 0 and 1 in ~ 12 h, *k*_p_ and *k*_dp_ should be of the order of 0.1 h^-1^.

#### ATPase reactions

Both CI and CII domains have ATPase activity, hydrolyzing about 10 ATP molecules in each CI domain and several ATP molecules in each CII domain in a day [23, 24]. It is reasonable to consider that ATP is consumed in the CII for supplying a phosphate group in the P process [40], but the reason for the ATP consumption in the CI has been elusive. With the present treatment *D*(*k,t*) implicitly represents the ATPase reactions in the CII, and we more focus on the ATPase reactions in the CI. We consider the case ATP is abundant in the solution; therefore, the probability that the CI binds no nucleotide is small. Hence, we use the variable 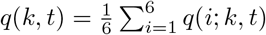 with

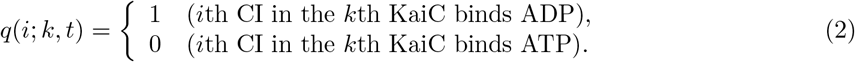

The ADP release and the subsequent ATP binding are the transition from *q*(*i*; *k,t*) = 1 to 0, and hydrolysis of the bound ATP is the transition from *q*(*i*; *k,t*) = 0 to 1. We simulate the stochastic ADP release and the ATP hydrolysis by treating *q*(*i*; *k,t*) as a stochastic variable changing with the lifetime of the ADP bound state, Δ_ADP_, and the frequency of hydrolysis, *f*_hyd_. The ATPase activity measured by the amount of the released ADP from KaiC is large in the P process [23], and the *X* ≈ 0 (*X* ≈ 1) structure binds ADP (ATP) [27]. These observations are consistent with the assumption that *f*_hyd_ is a constant independent of *X*(*k,t*) and Δ_ADP_(*k,t*) is a decreasing function of *X*(*k, t*). In practice, we represent this tendency as

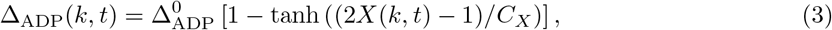

where 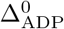 is a constant determining the timescale and *C_X_* is a constant determining the seisitivity to the structure.

#### Feedback coupling through structural change

Allosteric transitions in protein oligomers typically have a timescale of 10^-3^ ~ 10 ^−2^ s [63], and we assume a similar timescale in the present problem. Because the other reactions in our system are much slower, we describe the KaiC structure as in quasi-equilibrium by treating the chemical states as the quasi-static constraints. Representing the constraints with a variable *R*(*k,t*), we have the expression, 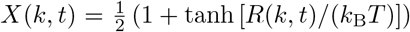 (See the Methods section). Here, the explicit form of *R*(*k, t*) constitutes major assumptions on the feedback couplings in the present model. Expanding *R*(*k,t*) up to the linear terms of *P*_C_6_A_2__, *P*_C_6_B_*i*_A_2*j*__, *D*(*k,t*), and *q*(*k,t*), we have

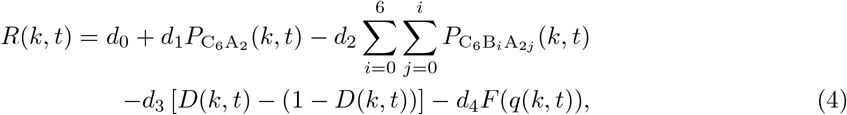

where *d*_0_ is a constant to determine the average structure, and *d*_1_, *d*_2_, *d*_3_, and *d*_4_ are constants defining the strength of the feedback coupling. We assume *d*_2_ > 0, which stabilizes the *X* ≈ 1 structure when KaiA binds on the CII, and *d*_2_ > 0, which stabilizes the *X* ≈ 0 structure when KaiB binds on the CI. Then, with the definitions of *h*_A_(*k,t*), *f*_A_(*k,t*), *h*_B_(*k,t*), and *f*_B_(*k,t*), we see that binding reactions constitute *the positive feedback loops* to stabilize the two states, the *X* ≈ 1 and *X* ≈ 0 states.

We use a constant *d*_3_ > 0 in Eq. 4, which destabilizes the *X* ≈ 1 state when phosphorylated (*D* ≈ 1) and destabilizes the *X* ≈ 0 state when dephosphorylated (*D* ≈ 0). This assumption is consistent with the X-ray crystallography analysis that Ser431 is phosphorylated in the *X* ≈ 0 structure while dephosphorylated in the *X* ≈ 1 structure [27]. Increase in *D* from *D* ≈ 0 to ≈ 1 is promoted by the KaiA binding to the CII of KaiC, which is promoted in the *X* ≈ 1 state, which then destabilizes the *X* ≈ 1 state through the *d*_3_ term. Decrease in *D* from *D* ≈ 1 to ≈ 0 is promoted by the KaiA unbinding from the CII of KaiC, which is promoted in the *X* ≈ 0 state, which then destabilizes the *X* ≈ 0 state through the *d*_3_ term. Therefore the *d*_3_ > 0 in Eq. 4 constitutes the negative feedback loops. Because *k*_p_ and *k*_dp_ in Eq. 1 are small, this destabilization of *X* is a slow process gradually proceeding after the structural transition. Therefore, the *d*_3_ term represents *the time-delayed negative feedback loops* to drive the oscillations between the *X* ≈ 1 and *X* ≈ 0 states.

*d*_4_*F*(*q*(*k,t*)) in Eq. 4 represents the coupling of structure with the ATPase reactions, which largely determines the balance between positive and negative feedback effects. This coupling was inferred from the observations that the ATP hydrolysis is necessary for binding KaiB to KaiC [51, 54, 64, 65], and that the structure is modified upon ATP hydrolysis [66, 24, 60, 27]. With *F*(*q*(*k,t*)) = *q*(*k,t*)*X* (*k, t*) – (1 – *q*(*k,t*))(1 – *X* (*k,t*)) and *d*_4_ > 0, the ADP binding on the CI destabilizes the *X* ≈ 1 state, while the ATP binding destabilizes the *X* ≈ 0 state. Therefore, the stochastic ATP hydrolysis triggers the transition to the *X* ≈ 0 state, and the stochastic release of ADP with the subsequent ATP binding triggers the transition to the *X* ≈ 1 state (Fig. 1).

Thus, the model describes the positive feedback between the structure and the binding/unbinding of KaiA and KaiB to/from KaiC (the *d*_1_ and *d*_2_ terms), the negative feedback between the structure and the P/dP process (the *d*_3_ term), and the transition-triggering effects of the ATPase reactions (the *d*_4_ term). These couplings generate the cooperative chemical and structural oscillations in individual KaiC molecules.

### Communication among many KaiC molecules at the ensemble level

We simulated the ensemble of *N* = 1000 or 2000 KaiC hexamers. For *N* = 1000 and *V* = 3 × 10^-15^l, the concentration of KaiC is *C*_T_ = 3.3 *μ*M on a monomer basis, which is near to the value 3.5 *μ*M often used in experiments. We assumed the ratio *A*_T_ : *B*_T_ : *C*_T_ = 1 : 3 : 3 as used in many experiments [67, 23, 54], where *A*_T_ and *B*_T_ are total concentrations of KaiA and KaiB on a monomer basis. The system was described by variables, *P*_C_6_A_2__(*k,t*), *P*_C_6_B_*i*_A_2*j*__(*k,t*), *D*(*k,t*), *X* (*k,t*), *q*(*k,t*), *x*_A_(*t*) and *x_B_*(*t*), with *k* = 1,…,*N*. *P*_C_6_B_*i*_A_2*j*__ (*k,t*) and *D*(*k,t*) were calculated by numerically integrating the kinetic equations, and *P*_C_6_A_2__(*k,t*) and *X*(*k,t*) were calculated with the quasi-equilibrium approximation at each time step. *q*(*k, t*) was calculated by simulating the stochastic transitions of *q*(*i*; *k,t*) between 0 and 1 with frequencies 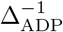 and *f*_hyd_. Concentrations of free unbound KaiA dimer and KaiB monomer, *x*_A_(*t*) and *x*_B_(*t*), were calculated at each time step from the following equations of conservation,

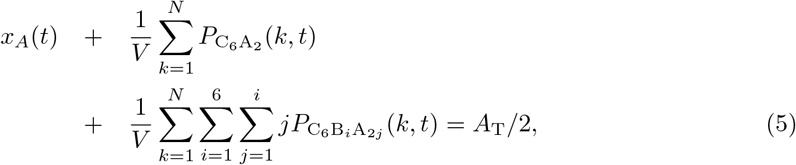

and 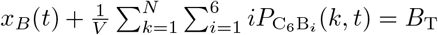. Competition between the 2nd and 3rd terms of the l.h.s. of Eq. 5 provides communication among KaiC molecules leading to the synchronization. See the Methods section for details.

## Results

### Single-molecule and ensemble-level oscillations

Figure 2 shows the calculated example oscillations. A KaiC hexamer arbitrarily chosen from the simulated ensemble of *N* = 1000 hexamers shows the structural transitions between the states *X*(*k, t*) ≈ 1 and *X*(*k, t*) ≈ 0 (Fig. 2A). The nucleotide-binding state in the CI ring *q*(*k, t*) also exhibits transitions between the state rapidly fluctuating around *q*(*k, t*) ≈ 0.3 and the state around *q*(*k, t*) ≈ 0.6. The phosphorylation level *D*(*k, t*) follows these switching transitions with the slower rates of the P/dP reactions; *D* increases in the *X* ≈ 1 state and decreases in the *X* ≈ 0 state, showing saw-tooth oscillations.

**Figure 2:**
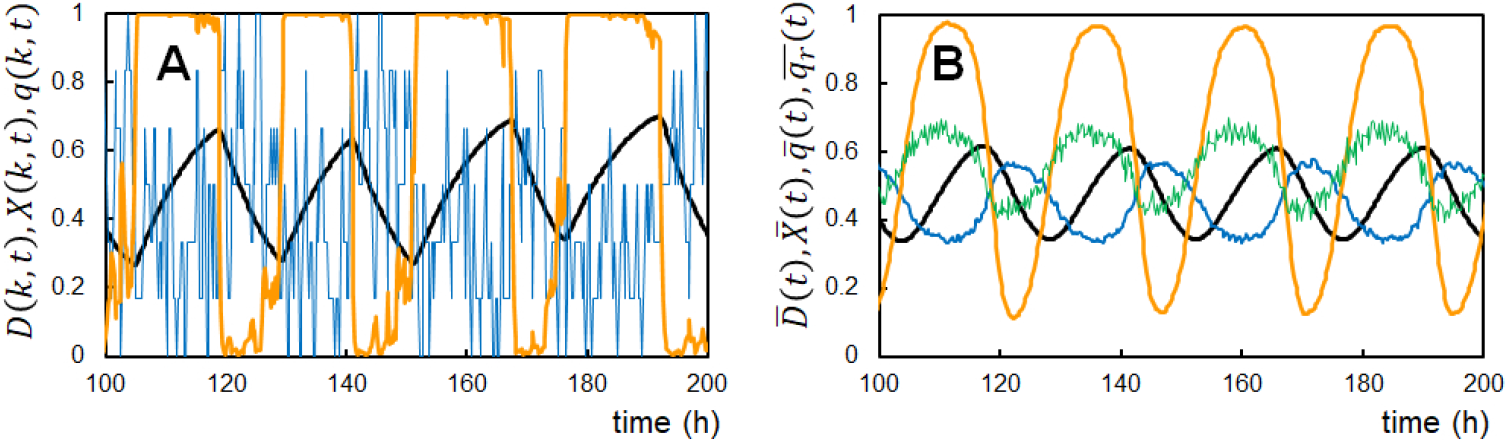
Example oscillations of the simulated KaiABC system. (**A**) The singlel-molecule oscillations of the phosphorylation level *D*(*k,t*) (black), the structural state *X* (*k,t*) (orange), and the ADP binding probability on the CI *q*(*k, t*) (blue) of an arbitrarily chosen *k*th KaiC hexamer. (**B**) The ensemble-averaged oscillations of the phosphorylation level 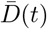 (black), the structural state 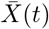 (orange), the binding probability of ADP on the CI 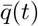 (blue), and the ATPase activity measured by the amount of the released ADP 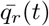 (green). *N* = 1000 at temperature *T*_0_ = 30 °C.

At the ensemble level, these fluctuating oscillations in individual molecules are averaged, resulting in the regular oscillations of 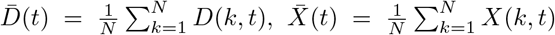, and 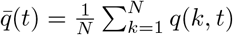 (Fig. 2B). The ensemble-averaged rate of the ADP release from KaiC, 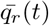 (Methods), is large during the P phase as observed experimentally [23], and the ADP binding probability 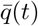 is large during the dP phase consistently with the experimental observations [27]. In this way, the model reproduces the stable circadian oscillations at the ensemble level averaging the synchronized individual KaiC oscillations.

### Modifications of the feedback strength

The binding of KaiA or KaiB may induce the global change of each KaiC subunit, shifting the position and orientation of subunit, whose effects represented by *d*_1_ and *d*_2_ in Eq. 4 should be insensitive to the single-residue substitution at the CI-CII interface. On the other hand, P/dP in the CII or the ATPase reaction in the CI is the local atomic change around the phosphate group, whose effects are transmitted through chains of electrostatic and volume-excluding interactions, producing the allosteric communication between the CI and CII [66, 27]. This process should be sensitive to the atomic interactions at the CI-CII interface and hence sensitive to the single-residue substitution. Here, we assume that substituting a CI-CII interface residue to the smaller volume one is represented by the decrease in *d*_3_ and *d*_4_ in Eq. 4 with a scaling factor *s* < 1 to *sd*_3_ and *sd*_4_ while *d*_4_ and *d*_2_ in Eq. 4 being kept in their original values. With *s* < 1, the simulated phosphorylation level oscillations indeed show the large amplitude and long period as expected from the decrease in the negative feedback strength, while the amplitude is small and the period is short for *s* > 1 (Fig. 3A). On the contrary, with positive feedback enhancement with a scaling factor *s* > 1, changing *d*_4_ and *d*_2_ to *sd*_4_ and *sd*_2_ enlarges the amplitude and period, while they are reduced when *s* < 1 (Fig. 3B).

**Figure 3:**
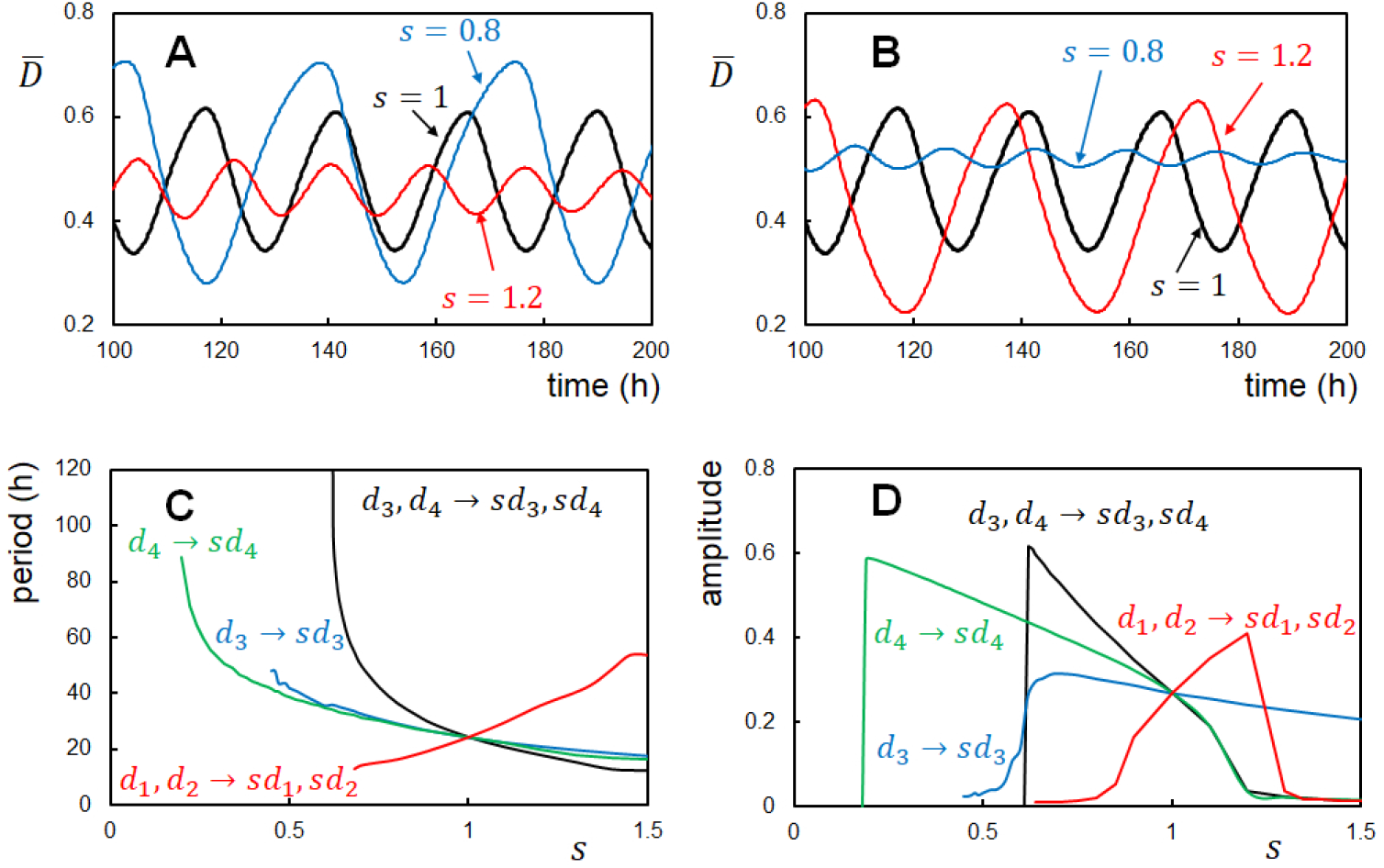
Effects of modification of the reaction-structure feedback strength. (**A**) and (**B**) Example oscillations of the ensemble-averaged phosphorylation level 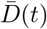 with a varying factor *s*. (**A**) The negative feedback coupling between structure and reactions was modified; the coupling with the P/dP reactions *d*_3_ and the one with the ATPase reactions *d*_4_ were modified to *sd*_3_ and *sd*_4_ with *s* = 0.8 (blue), 1.0 (black), and 1.2 (red). (**B**) The positive feedback coupling between structure and the binding/unbinding reactions was modified; the coupling with the KaiA binding/unbinding *d*_1_ and the one with the KaiB binding/unbinding *d*_2_ were modified to *sd*_1_ and *sd*_2_ with *s* = 0.8 (blue), 1.0 (black), and 1.2 (red). Period (**C**) and amplitude (**D**) of the simulated 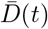 oscillations plotted as functions of the scaling factor s for modifying *d*_3_ to *sd*_3_ while *d*_1_, *d*_2_, and *d*_4_ being kept constant (blue), *d*_4_ to *sd*_4_ while *d*_1_, *d*_2_, and *d*_3_ being kept constant (green), *d*_3_ and *d*_4_ to *sd*_3_ and *sd*_4_ while *d*_1_ and *d*_2_ being kept constant (black), and *d*_1_ and *d*_2_ to *sd*_1_ and *sd*_2_ while *d*_3_ and *d*_4_ being kept constant (red). *N* = 1000 and temperature was *T*_0_ = 30 °C.

These effects of the feedback strength modifications were systematically examined in Figs. 3C and 3D. When *d*_3_ is scaled to *sd*_3_ with *d*_4_, *d*_4_, and *d*_2_ kept constant, the amplitude and period are extended as *s* is decreased. When *s* < 0.5, the oscillations disappear as the system is caught at the *X* ≈ 1 state losing the negative feedback destabilization of the *X* ≈ 1 state. Similar behaviors were found when the structure-ATPase coupling was changed to *sd*_4_ with *d*_3_, *d*_4_, and *d*_2_ kept constant. Thus, the ATPase reactions give similar effects to the negative feedback in the present model. With the combined change to *sd*_3_ and *sd*_4_, the period change is more prominent, ranging from 9 h to 120 h (Fig. 3C), which explains the observed ten-times period change induced by the single-residue substitution at the CI-CII interface [26]. Modifying the positive feedback strength to *sd*_4_ and *sd*_2_ shows that the oscillations disappear when the positive feedback is too weak or too strong (Fig. 3D). The period and amplitude are enlarged as s increases in between these boundaries of positive feedback strength (Fig. 3C).

### Thermal loosening of the strcutural coupling explains temperature compensation

We propose that the increased structural fluctuations of KaiC in the higher temperature should weaken the interactions at the CI-CII interface to decrease *d*_3_ and *d*_4_ in Eq. 4, producing similar effects to the single-residue substitution (Fig. 3A). These effects should enlarge the amplitude and period, compensating for the accelerated reactions in high temperatures. Here, we examine our hypothesis by simulating the oscillations in various temperatures.

Assuming that the fluctuations are proportinal to exp(-Δ*E*_f_/(*k*_B_*T*)) with a constant Δ*E*_f_, the feedback coupling strength at temperature *T*, *d*_3_(*T*) and *d*_4_(*T*), should be modified from those in the standard temperature *T*_0_ = 30°C as *d*_3_(*T*) = *d*_3_(*T*_0_)/*s*(*E*_f_;*T*, *T*_0_) and *d*_4_(*T*) = *d*_4_(*T*_0_)/*s*(*E*_f_; *T*, *T*_0_);

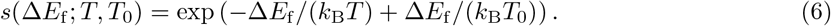

We call this temperature dependence of *d*_3_(*T*) and *d*_4_(*T*) Rule 1. Another rule comes from the activation energy Δ*E*_gs→fs_ > 0 and Δ*E*_fs→gs_ > 0 for the KaiB fold transformations. We assume *h*_B_ ∝ exp [–Δ*E*_gs→fs_/(*k*_B_*T*)] and 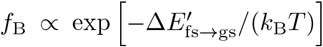, which affects the binding/unbinding kinetics of KaiB. Here, 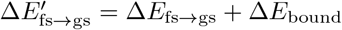 and Δ*E*_bound_ is the KaiC-KaiB binding energy. We call this assumption on *h*_B_ and *f*_B_ Rule 2. The other assumption corresponds to the temperature insensitivity of the ATPase reactions as observed in experiments [23, 68]. We assume the temperature-insensitive ATPase reactions by imposing 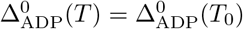 and *f*_hyd_(*T*) = *f*_hyd_(*T*_0_), where 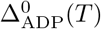 is a constant to determine the lifetime of the ADP bound state (Eq. 3), and *f*_hyd_(*T*) is the hydrolysis frequency of the bound ATP. We call this assumption Rule 3. These three rules are summarized in Table 1.

**Table 1:** Temperature dependence/independence of specific parameters.

| Rules | $T$-dependence* | Physical implications |
| --- | --- | --- |
| Rule 1 | $d_3(T) = \frac{d_3(T_0)}{s(\Delta E_f; T, T_0)}$ | Thermally attenuated<br>structure-P/dP feedback |
| | $d_4(T) = \frac{d_4(T_0)}{s(\Delta E_f; T, T_0)}$ | Thermally attenuated<br>structure-ATPase coupling |
| Rule 2 | $h_B(T) \propto s(\Delta E_0; T, T_0)$<br>$\times s(\Delta E_{gs \rightarrow fs}; T, T_0)$ | Thermally activated<br>KaiB fold |
| | $f_B(T) \propto s(\Delta E_0; T, T_0)$<br>$\times s(\Delta E'_{fs \rightarrow gs}; T, T_0)$ | |
| Rule 3 | $\Delta_{ADP}^0(T) = \Delta_{ADP}^0(T_0)$ | $T$-insensitive lifetime<br>of the ADP bound state |
| | $f_{hyd}(T) = f_{hyd}(T_0)$ | $T$-insensitive frequency<br>of ATP hydrolysis |
| Case A | Rule 1+Rule 2+Rule 3 |  |
| Case B | Rule 1+Rule 3 | $\Delta E_{gs \rightarrow fs} = \Delta E'_{fs \rightarrow gs} = 0$ |
| Case C | Rule 1+Rule 2 | $\Delta_{ADP}^0(T)^{-1} \propto s(\Delta E_0; T, T_0)$<br>$f_{hyd}(T) \propto s(\Delta E_0; T, T_0)$ |
| Case D | Rule 2+Rule 3 | $\Delta E_f = 0$ |
| *$T_0 = 30^\circ\text{C}$, $\Delta E_0 = 10k_B T_0$, $\Delta E_f = 6-7k_B T_0$,<br>$\Delta E_{gs \rightarrow fs} = 10k_B T_0$, and $\Delta E'_{fs \rightarrow gs} = 12k_B T_0$. | | |

The model has six rate constants (Table 2) other than the four rate constants, *h*_B_, *f*_B_, 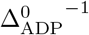, and *f*_hyd_, discussed in Table 1. Each of six rate constants should depend on temperature in each distinctive way. However, for examining the hypothesis transparently, we assume a straightforward case of the same activation energy Δ*E*_0_ in six rate constants, which leads to the same temperature dependence as *k*_dp_(*T*) = *s*(Δ*E*_0_; *T, T*_0_)*k*_dp_(*T*_0_), etc., where s is defined in Eq. 6. In the case we do not impose any of Rule 1, Rule 2, or Rule 3 by assuming 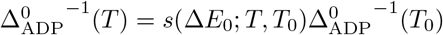 and *f*_hyd_(*T*) = *s*(Δ*E*_0_; *T, T*_0_)*f*_hyd_(*T*_0_), all 10 rate constants are scaled in the same way, which is almost the same as the scaling in time units, making the oscillation period approximately proportional to *s*(Δ*E*_0_; *T, T*_0_)^-1^. This homogeneous activation without thermal attenuation of feedback should result in *Q*_10_ = (period in *T*_0_–5 °C)/(period in *T*_0_+5 °C) ≈ *s*(Δ*E*_0_; *T*_0_+5, *T*_0_–5) ≈ 1.4 for Δ*E*_0_ = 10*k*_B_*T*_0_ and *Q*_10_ ≈ 2 for Δ*E*_0_ = 20*k*_B_*T*_0_. The purpose of the present subsection is to show that |*Q*_10_ – 1| ≲ 0.1, or the oscillations are temperature compensated when we assume the three rules in Table 1 even in the case Δ*E*_0_ = 10*k*_B_*T*_0_ in other six rate constants. We also show that Rule 1 plays a dominant role, and temperature compensation is realized without imposing Rule 2 or 3; temperature compensation is realized if Rule 1 is satisfied even when Δ*E*_0_ = 10*k*_B_*T*_0_ is used for all ten rate constants.

**Table 2:**
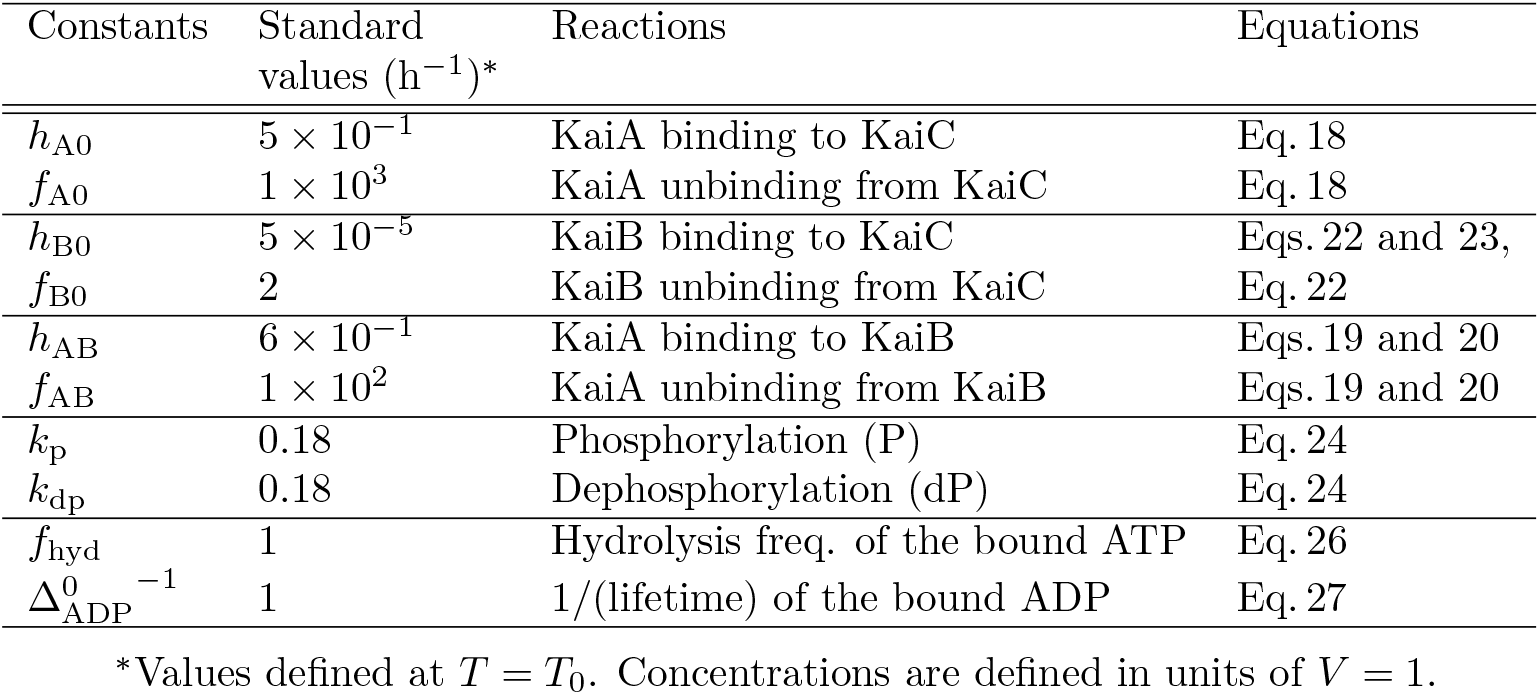
Rate constants in the model.

In order to distinguish the roles of three rules, we examined four cases (Table 1). All three Rules apply in Case A (Fig. 4A), Rules 1 and 3 in Case B (Fig. 4B), Rules 1 and 2 in Case C (Fig. 4C), and Rules 2 and 3 in Case D (Fig. 4D). The period is temperature compensated in Case A (*Q*_10_ = 1.01), Case B (*Q*_10_ = 0.93), and Case C (*Q*_10_ = 1.02) with a slight overcompensation in Case B, while the period shows a distinct temperature dependence in Case D (*Q*_10_ = 1.51), showing that Rule 1 plays a dominant role in temperature compensation. The overcompensation in Case B shows that Rule 2 enhances the temperature dependence of the period, which was compensated by Rule 1 in Cases A and C. In Cases A, B, and C, amplitude sharply decreases as temperature decreases below 20 °C as observed experimentally [69]. We should note that Rule 3, temperature insensitivity of ATPase reactions, does not play a significant role in the present examination (Fig.4C); we discuss more on this point in the “Effects of the ATPase reactions” subsection.

**Figure 4:**
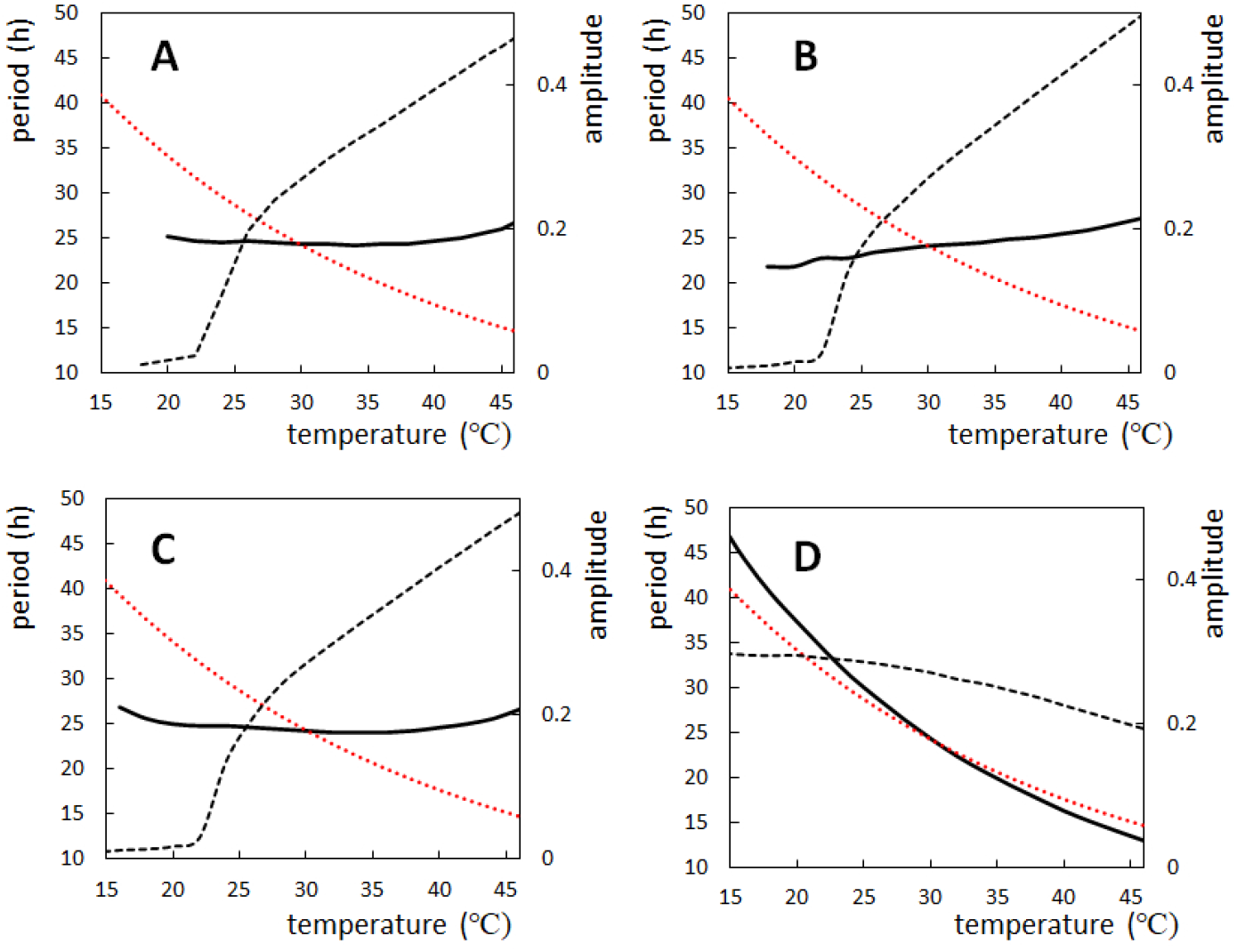
Temperature dependence of the period and amplitude of the simulated oscillations. Period (solid line) and amplitude (dashed line) of the ensemble-averaged phosphorylation level 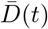 were calculated as functions of temperature with various modeling rules. Compared is the relative change of 1/rate of the process having the 10*k*_B_*T*_0_ activation energy (red dotted line). (**A**) Rule 1 (thermal attenuation of the negative feedback coupling), Rule 2 (thermal transformation of KaiB to a bindable conformation), and Rule 3 (temperature-insensitive ATPase reactions) applied (Case A). (**B**) Rule 1 and Rule 3 applied (Case B). (**C**) Rule 1 and Rule 2 applied (Case C). (**D**) Rule 2 and Rule 3 applied (Case D). *N* = 2000.

A determinant role of Rule 1 is evident also from calculations with the varied temperature dependence of the structural fluctuations. *Q*_10_ decreases as Δ*E*_f_ increases (Fig. 5A); *Q*_10_ = 1.51 (Δ*E*_f_ = 0), 1.28 (Δ*E*_f_ = 3*k*_B_*T*_0_), 1.08 (Δ*E*_f_ = 6*k*_B_*T*_0_), and 0.90 (Δ*E*_f_ = 9*k*_B_*T*_0_); showing the temperature compensation with Δ*E*_f_ = 6*k*_B_*T*_0_, and the distinct overcompensation with 9*k*_B_*T*_0_. A drop of the amplitude in the low temperature regime takes place at the higher temperature as ΔEf increases (Fig. 5B), consistently with the expected period-amplitude correlation.

**Figure 5:**
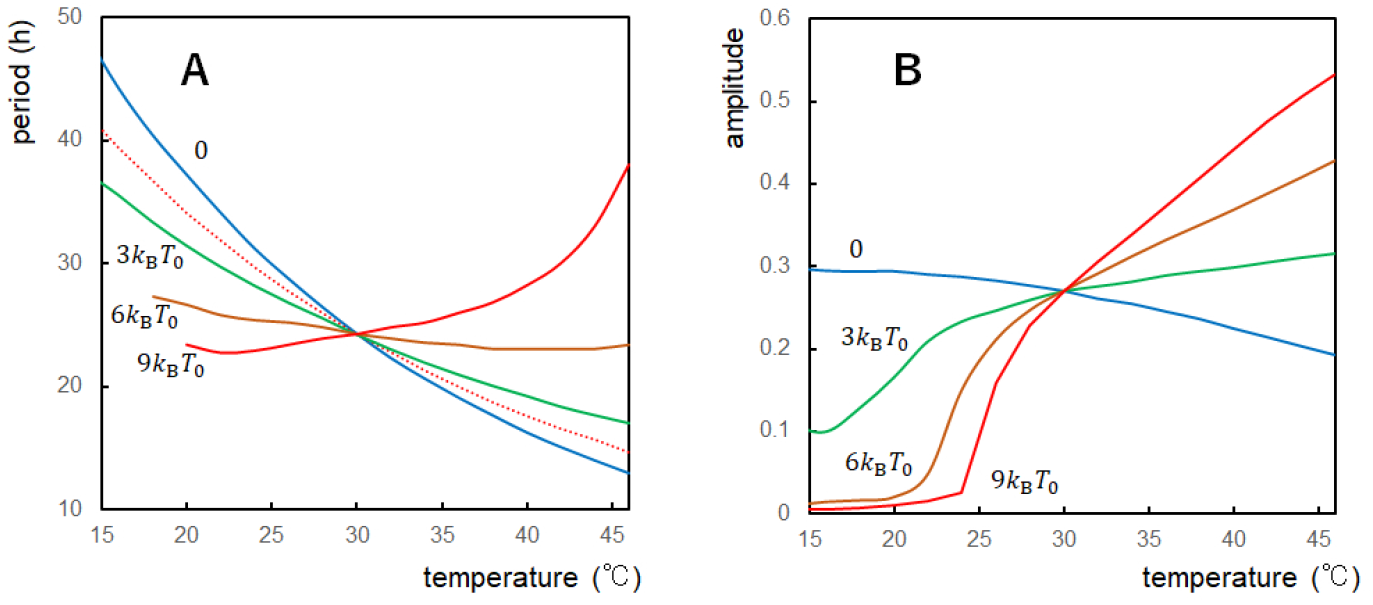
Temperature dependence of the period and amplitude simulated with a different activation energy of structural fluctuations. Period and amplitude of the ensemble-averaged phosphorylation level 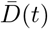 were calculated with various values of the activation energy of structural fluctuations Δ*E*_f_ with Δ*E*_f_ = 0 (blue), 3*k*_B_*T*_0_ (green), 6*k*_B_*T*_0_ (brown), and 9*k*_B_*T*_0_ (red) with *T*_0_ = 30 °C. (**A**) Period plotted for each Δ*E*_f_ as a function of temperature. Compared is the relative change of 1/rate of the process having the 10kBT0 activation energy (red dotted line). (**B**) Amplitude plotted for each Δ*E*_f_ as a function of temperature. *N* = 2000.

Fig. 5 shows a sensitive dependence of *Q*_10_ on Δ*E*_f_. In KaiC mutants showing the altered oscillation period, we can expect the intact *Q*_10_ ≈ 1 when the mutations perturb the P/dP-reaction rates in the CII or the ATPase-reaction rates in the CI, which alters the period, but does not alter the CI-CII interface, and hence does not affect Δ*E*_f_ much. However, with a single-residue substitution at the CI-CII interface of KaiC, the substitution can alter Δ*E*_f_. In the experimental report, some substitutions at the CI-CII interface showed the intact *Q*_10_, but the others showed the enlarged *Q*_10_ [26]. A possible explanation for the former case is that the protein region around the substituted residue compensates for the structural fluctuations’ temperature dependence, leading to the robust Δ*E*_f_ against the mutation, while in the latter case, such compensation does not sufficiently work, leading to the enlarged *Q*_10_. It is crucial to examine whether such a difference in Δ*E*_f_ indeed occurs in those mutants.

### Phase shift caused by a stepping change in temperature

A significant feature of circadian rhythms is the response of their oscillation phase to the external stimuli. In particular, the phase is distinctively shifted by a temperature change in circadian oscillators while their period is temperature compensated [70, 71]. Also, in the in vitro KaiABC system, Yoshida et al. [72] found the phase shift indued upon temperature change. A step-up of temperature from *T* = 30 ^°^C to 45 ^°^C in the dP process advanced the phase, while a step-up in the P process delayed the phase. On the other hand, a step-down of temperature from *T* = 45 ^°^C to 30 ^°^ C in the P process advanced the phase, and a step-down in the dP process delayed the phase. Here, we compare the simulated phase shift in the present model with the experimental results in [72]. In order to avoid the confusion coming from the slight period change upon temperature stepping, we defined the circadian time (CT) in the same way as in Ref.[72]: timing of the peak in the simulated rhythm at each temperature was set to CT 16, and the period length was normalized to be 24 h in CT (Fig. 6A).

**Figure 6:**
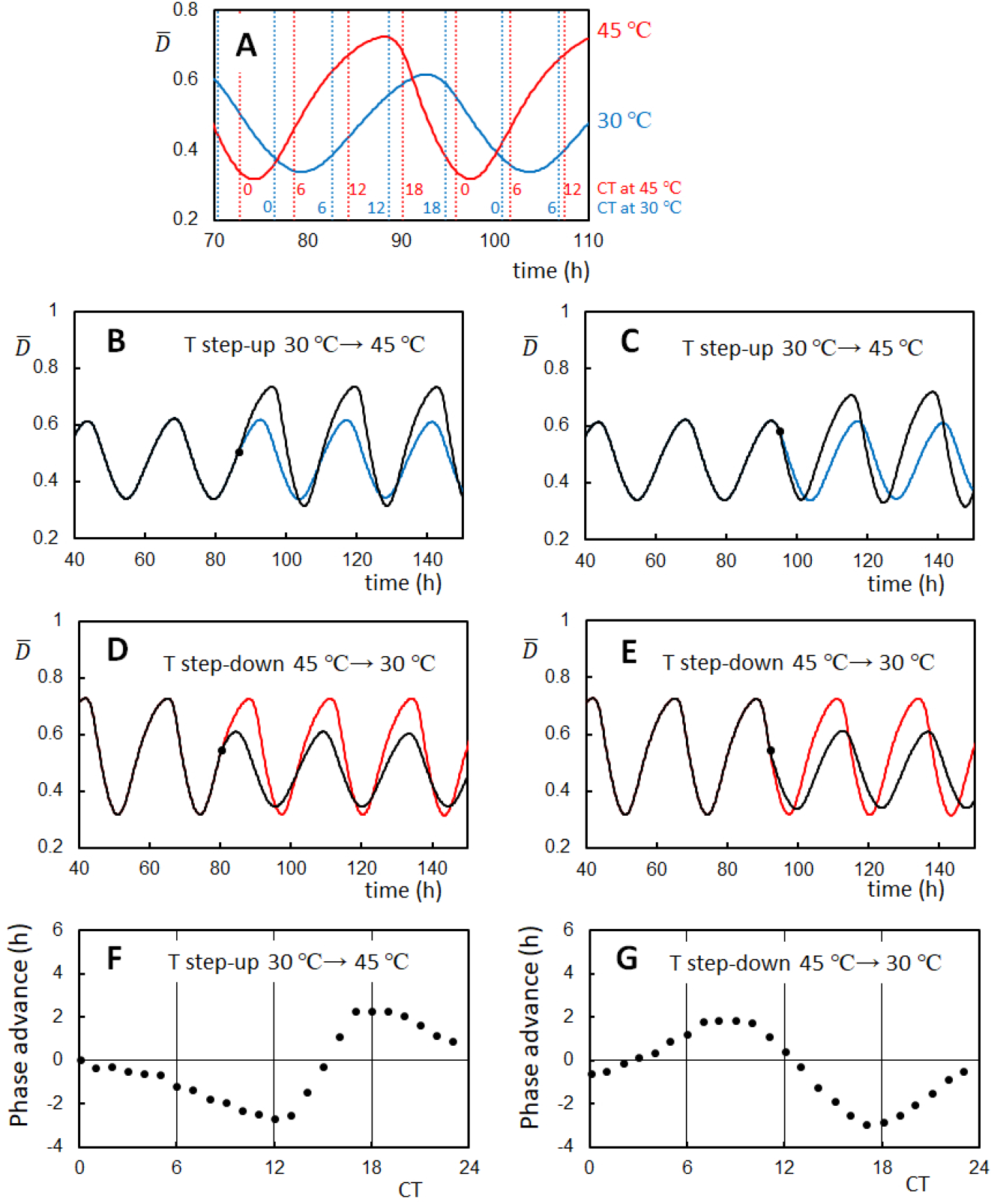
Phase shifts of the ensemble-averaged oscillations of the KaiC phosphorylation level caused by temperature steps. (**A**) Circadian Time (CT) was set to be 16 at the peak of oscillations at each temperature and the period was normalized to be 24 h in CT. Time denoted on the x-axis is the simulated incubation time (IT) (**B**) An example trajectory showing the phase delay upon temperature step-up from *T* = 30 °C to *T* = 45°C at CT 9 (filled circle at IT 85.6 h). (**C**) An example trajectory showing the phase advance upon temperature step-up from *T* = 30 °C to *T* = 45 °C at CT 17.3 (filled circle at IT 95). In **B** and **C**, The oscillation trajectory going through the temperature step-up (black) and the trajectory with the constant temperature at *T* = 30 °C (blue) are superposed. (**D**) An example trajectory showing the phase delay upon temperature step-down from *T* = 45°C to *T* = 30°C at CT 7.9 (filled circle at IT 80.4). (**E**) An example trajectory showing the phase advance upon temperature step-down from *T* = 45°C to *T* = 30 °C at CT 20 (filled circle at IT 92). In **D** and **E**, The oscillation trajectory going through the temperature step-down (black) and the trajectory with the constant temperature at *T* = 45°C (red) are superposed. (**F**) Phase response curve (PRC) plotted as a function of CT when the temperature was stepped up from *T* = 30°C to *T* = 45 °C. (**G**) PRC plotted as a function of CT when the temperature was stepped down from *T* = 45°C to *T* = 30°C. Rule 1, Rule 2, and Rule 3 in Table 1 were used with Δ*E*_f_ = 6*k*_B_*T*_0_ and *N* = 1000.

In the present model, oscillations of individual KaiC molecules are synchronized by the KaiA binding to the KaiC-KaiB-KaiA complexes, which entrains oscillating molecules in the ensemble into the dP phase [44]. With Rule 2 in Table 1, the *T* step-down increases the ratio *h*_B_/*f*_B_, enhancing the entrainment and stabilizing the dP phase. Thus, the *T* step-down in the dP phase elongates the dP process and causes the phase delay. On the other hand, the *T* step-up in the same dP phase causes the opposite effect of the phase advance. This expectation is confirmed in the results shown in Figs. 6B–6G. In Figs. 6B and 6C, the temperature was stepped up from *T* = 30 °C to 45 °C. Fig. 6B shows the phase delay caused by a simulated *T* step-up in the P phase at CT 9 (85.6h in the simulated incubation time (IT)), and Fig. 6C shows the phase advance caused by a *T* step-up in the dP phase at CT 17.3 (IT 95). In Figs. 6D and 6E, the temperature was stepped down from *T* = 45 °C to 30 °C. Fig. 6D shows the phase advance caused by a simulated *T* step-down in the P phase at CT 7.9 (IT 80.4), and Fig. 6E shows the phase delay caused by a *T* step-down in the dP phase at CT 20 (IT 92). These results can be quantified by plotting a phase response curve (PRC). Fig. 6F and Fig. 6G show PRCs for *T* step-up and *T* step-down, respectively. These simulated PRCs qualitatively reproduce the experimentally observed PRCs in [72]: upon *T* step-up, the phase was advanced at CT 16–CT 22 and delayed at CT 8–CT 14, and upon *T* step-down, the phase was advanced at CT 6–CT 11 and delayed at CT 14–CT 24. Thus, our model reproduces the essential features of the experimentally observed phase shift upon the stepping change of temperature.

### Effects of the ATPase reactions

In order to clarify the effects of the ATPase reactions on the oscillations, we scaled the rate constants of the ATPase reactions in the present model. Fig. 7A shows the amplitude and period calculated by modifying the inverse lifetime of the ADP bound state, 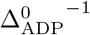, and the hydrolysis frequency of the bound ATP, *f*_hyd_. They were scaled by a factor *s*_a_ in two different ways; in case I, they were scaled as 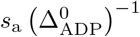 and *s*_a_*f*_hyd_, and in case II, 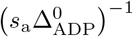 and *s*_a_*f*_hyd_. In case I, period and amplitude only slightly depend on *s*_a_ for *s*_a_ > 0.5. This insensitivity to the scaling corresponds to the temperature insensitivity shown in Fig. 4C, where we did not use Rule 3 but scaled the constants as *s*(Δ*E*_0_; *T, T*_0_) 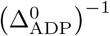 and *s*(Δ*E*_0_; *T*, *T*_0_)*f*_hyd_. The insensitivity in case I and Fig. 4C indicates that the change in the ATPase reaction rates does not affect the period when they were changed with the constraint Δ_ADP_ · *f*_hyd_ = const. The ATPase reactions should affect the period in two ways: The probability that the CI binds ADP during the P phase is correlated to how the *X* ≈ 1 state is destabilized, and the probability that the CI binds ATP during the dP phase is correlated to how the *X* ≈ 0 state is destabilized. However, changing the rates under the constraint Δ_ADP_ · *f*_hyd_ = const. does not affect these probabilities. Therefore, the period was insensitive to the changes in the rate constants under the constraint Δ_ADP_ · *f*_hyd_ = const.

**Figure 7:**
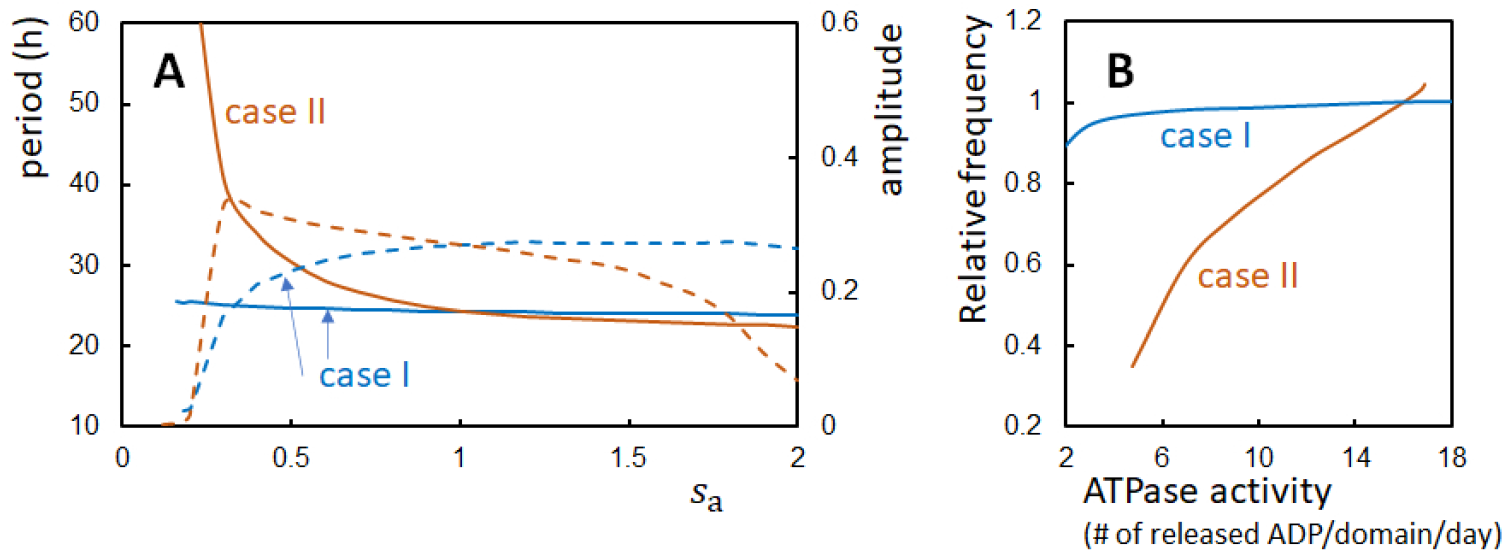
Oscillations and the ATPase activity calculated with the modified rate constants of the ATPase reactions. Two ways of the modification, case I and case II, were tested on the inverse lifetime of the ADP bound state, 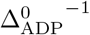, and the frequency of hydrolysis of the bound ATP, *f*_hyd_. In case I (blue), the rate constants were scaled as 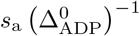 and *s*_a_*f*_hyd_, and in case II (brown), 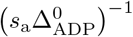 and *s*_a_*f*_hyd_ (**A**) The period (solid line) and amplitude (dashed line) of the ensemble-averaged phosphorylation level 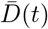 were calculated as functions of *s*_a_. (**B**) The frequency (inverse period) of oscillations of 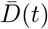 is compared with the ATPase activity (in units of the number of released ADP molecules from a CI domain in a day). The ATPase activity was calculated in the nonoscillatory condition in the absence of KaiA and KaiB. Temperature is *T*_0_ and *N* = 2000.

In case II, this constraint is not satisfied, and the change in the rate constants brings about a significant change in the period (Fig. 7A). We examined the correlation between the oscillation frequency (i.e., 1/period) and the ATPase activity calculated in the nonoscillatory condition in the absence of KaiA and KaiB. Fig. 7B shows a clear correlation between the oscillation frequency and the ATPase activity in case II as experimentally observed [23, 24]. The period is insensitive to the change in the ATPase rates only when the specific constraint is satisfied. This result suggests that the strategy to keep the specific constraint has been avoided in evolutionary design. Instead, the more general strategy of the temperature-insensitive ATPase reactions might have been evolutionarily realized in KaiC [23, 68] as in the TTO regulator, CKI*ε*/*δ*, in mammals [22].

### Phase shift caused by a pulse of the increased ADP concentration

Examining the response of the oscillation phase to the change in the ATP/ADP concentration ratio should further test the roles of the ATP hydrolyses in the KaiABC oscillations. Rust et al. [73] experimentally examined the phase response of the in vitro KaiABC system by adding ADP to the reaction buffer at various timing. After several hours of ADP addition, they reduced the ADP concentration to the original level by adding pyruvate kinase, which converts ADP to ATP. Rust et al. found that this pulse of ADP addition delayed the oscillation phase when the pulse was added at around the trough of the oscillating phosphorylation level, while the pulse advanced the phase when the pulse was added at around the peak of the phosphorylation level [73]. In our model, the increased concentration of ADP should elongate the ADP-bound state’s lifetime, enhancing the transition probability from the P phase to the dP phase and reducing the probability of the opposite transition. Therefore, we expect that adding the ADP pulse at the oscillation peak helps the transition to advance the phase, while adding the ADP pulse at the oscillation trough disturbs the transition to delay the phase. This expectation is consistent with the results observed in [73], and we confirmed it with simulations as in the following.

In our model, the effects of an increase in the ADP concentration should correspond to the increase in the ADP-bound state’s lifetime. Therefore, we simulated the ADP pulse by increasing 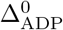 at a particular time and reducing 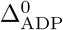 to the original value 6 hours after the increase. We defined CT in the same way as in Fig. 6 (Fig. 8A) and used the same parameters as in Fig. 4A. Fig. 8B and Fig. 8D show the example phase shift with the simulated addition of ADP pulse around the trough and the peak of oscillations, respectively. As expected, the addition of the pulse around the trough delayed the oscillation phase (Fig. 8B), and the addition of the pulse around the peak advanced the phase (Fig. 8D). It is intriguing that the simulated change from the phase-delay to phase-advancement behaviors was abrupt, inducing a jump in the PRC at around CT 5 (Fig. 8E), reproducing the observed jump in the experimental PRC [73]. The model predicts that the addition of the ADP pulse at the jump point at CT 5 (IT 81.6) advanced the oscillation phase of almost half of KaiC molecules and delayed the phase of the rest half of molecules, which strongly desynchronized the oscillations of individual KaiC molecules, diminishing the ensemble-level oscillations (Fig. 8C); it is important to experimentally examine this desynchronization as a test of the oscillation mechanism proposed in the present model.

**Figure 8:**
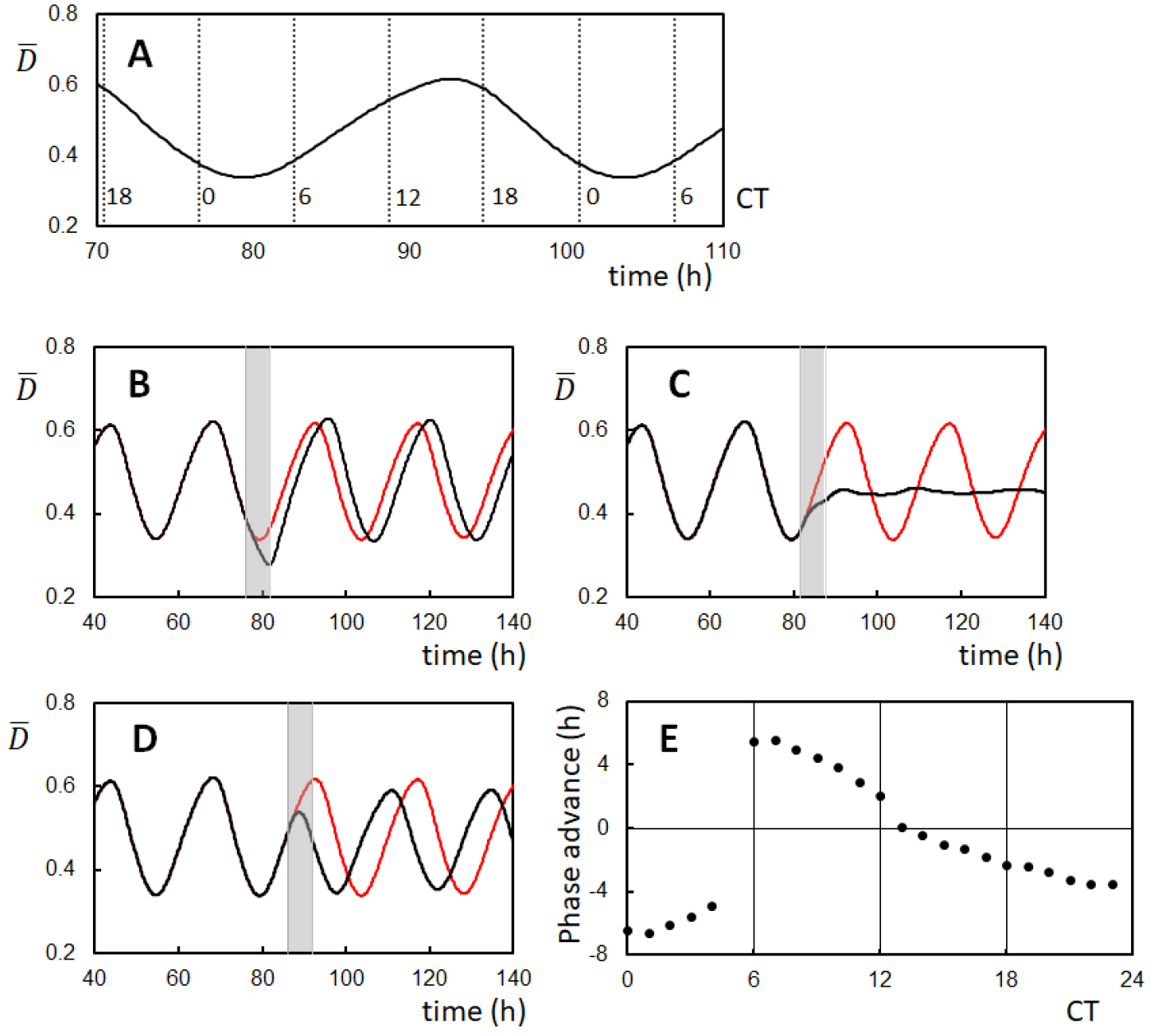
Phase shifts of the ensemble-averaged oscillations of the KaiC phosphorylation level caused by pulses of the ADP increase. (**A**) Circadian Time (CT) was set to be 16 at the peak of oscillations and the period was normalized to be 24 h in CT. Time denoted on the x-axis is the simulated incubation time (IT). (**B**) An example trajectory showing the phase delay by adding an ADP pulse from CT 0 to CT 6 (gray bar from IT 76 to IT 82). (**C**) An example trajectory showing the desynchronization by adding an ADP pulse from CT 5 to CT 11 (gray bar from IT 81.6 to IT 87.6). (**D**) An example trajectory showing the phase advance by adding an ADP pulse from CT 9.5 to CT 15.5 (gray bar from IT 86 to IT 92). In **B**, **C**, and **D**, the oscillation trajectory perturbed by an ADP pulse (black) and the trajectory without perturbation (red) are superposed. (**E**) PRC plotted as a function of CT when the ADP pulse started to be added. During the ADP pulse, 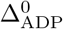 was increase to 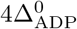. Rule 1, Rule 2, and Rule 3 in Table 1 were used with *T* = *T*_0_ and *N* = 1000.

## Discussion

Using the model describing the feedback coupling among reactions and structural transitions in individual KaiC molecules and synchronization of many KaiC molecules, we showed that weakening the negative feedback strength extends the amplitude and lengthens the period of oscillations in the KaiABC system. This weakening can explain the observed wide range of period modification induced by the single-residue substitution at the CI-CII interface of KaiC [26]. We hypothesized that thermal fluctuations induce similar effects to the substitution at the CI-CII interface, which explained the stable temperature compensation in the KaiABC system. The ATPase reactions also affect the period, but the period is insensitive when the ATPase rates are changed under the specific constraint.

A possible test of the thermal weakening of the CI-CII structural coupling is to measure the temperature dependence of the thermal fluctuations of the interface residue with NMR or other spectroscopic methods. The model predicts that the temperature dependence of fluctuations should be anti-correlated with the *Q*_10_ of the period, and such an anti-correlation was suggested from the recent neutron scattering data [68].

Another possible test is to observe the single-molecular behavior. In the typical oscillatory condition, oscillations of individual KaiC molecules are synchronized, inducing coherent circadian oscillations at the ensemble level (Fig. 9A). If the synchronization is realized through the sequestration of KaiA into the KaiC-KaiB-KaiA complexes as assumed in the present study, synchronization should be lost when the binding affinity between KaiA and KaiB is reduced. It would be possible to design a KaiB mutant with a low affinity to KaiA, and in the present model, such a mutant is represented by reducing the *h*_AB_ value. With this reduction, synchronization is lost, and the ensemble oscillations disappear despite the oscillations with the large amplitude remaining in individual KaiC molecules (Fig. 9B). In such a desynchronized condition, period of individual oscillations is temperature compensated with the temperature-dependent oscillation amplitude in the present model (Fig. 9C). In contrast, when the negative feedback strength does not depend on temperature, individual oscillations are not temperature compensated without showing a significant temperature dependence of amplitude (Fig. 9D). The temperature compensation hypothesis in the present study contrasts with the hypothesis of the competitive KaiA binding reactions [25]. With the latter hypothesis, temperature compensation is induced only from the ensemble level mechanism, not from the individual molecular mechanism, which should be distinguishable with the single-molecule observation in the condition that the synchronization is lost.

**Figure 9:**
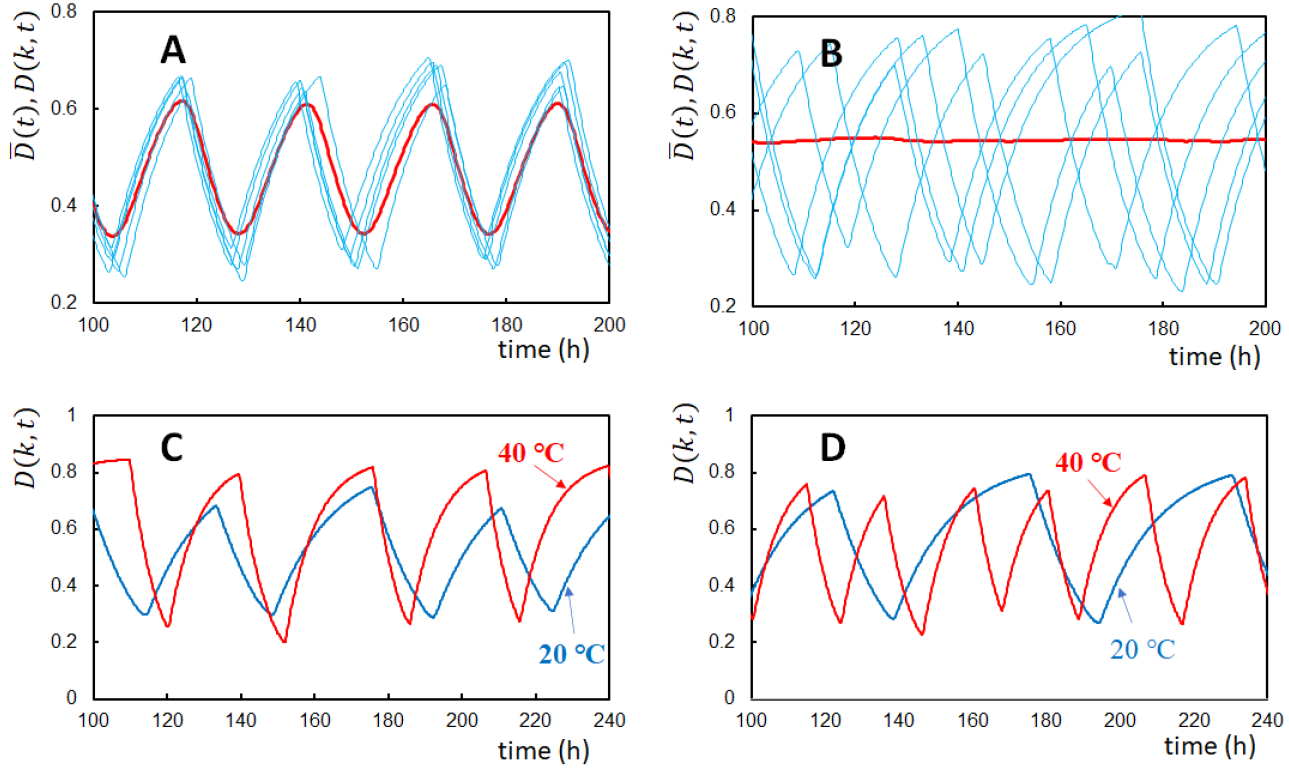
Temperature compensation of oscillations of individual molecules. In **A** and **B**, oscillations of the ensemble-averaged phosphorylation level 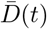 (red) are compared with five individual oscillations *D*(*k,t*) of molecules arbitrarily chosen from the ensemble (blue). *T* = *T*_0_. (**A**) *h*_AB_ was kept in the standard value (Table 2). (**B**) *h*_AB_ was reduced to 1/20 of the standard value. (**C**) Oscillations of *D*(*k,t*) of an arbitrarily chosen single molecule at *T* = 40°C (red) and *T* = 20°C (blue). Three rules, Rule 1 (thermal attenuation of the reaction-structure feedback coupling with Δ*E*_f_ = 6*k*_B_*T*), Rule 2 (thermal activation of KaiB transformation), and Rule 3 (temperature insensitivity of the ATPase reactions), were assumed. (**D**) Oscillations of *D*(*k, t*) of an arbitrarily chosen single molecule at *T* = 40 °C (red) and *T* = 20 °C (blue). Rule 2 and Rule 3 were assumed, while Rule 1 was not adopted with Δ*E*_f_ =0. In **C** and **D**, *h*_AB_ was reduced to 1/20 of the standard value. *N* = 1000.

With the larger KaiA concentration than the standard value of *A*_T_/*C*_T_ = 1/3, Ito-Miwa et al. showed that *Q*_10_ of the oscillation period becomes large (Fig. S6 in [26]). In our model with the parameters used in Fig. 6, with the increased KaiA concentration of *A*_T_/*C*_T_ = 1/2, the oscillation period was temperature compensated with *Q*_10_ = 0.98, which disagrees with the experimental report. We should note that the results depend on the parameterization in the model; for example, with Δ*E*_gs→fs_ = 20*k*_B_*T* and 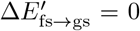, *Q*_10_ = 1.33 when *A*_T_/*C*_T_ = 1/2. Therefore, the further examinations should be needed to compare the model results with the experimental report by carefully examining the mechanisms of the binding/unbinding kinetics of KaiA and KaiB to/from KaiC. Our analyses showed the importance of the molecular features which determine the coupling between reactions and structural transitions in the oscillating molecules. Investigations of the KaiABC system should highlight the significance of the regulation through the atomic reaction-structure coupling and help provide innovative methods of design for regulating the system dynamics of an ensemble of molecules.

## Methods

### Multifold feedback coupling in KaiC

We describe individual KaiC hexamers with coarse-grained variables; the binding state of KaiA on the CII domain, *θ*_C_6_A_2__(*k*), the binding state of KaiB and KaiA on the CI domain, *θ*_C_6_B_*i*_A_2*j*__(*k*), the phosphorylation level, 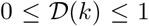, the structural state, *W*(*k*), and the nucleotide-binding state, *q*(*i*; *k,t*). Here,

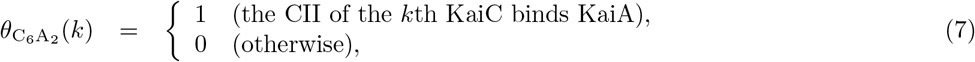

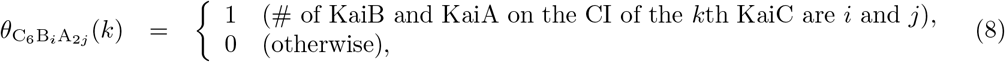

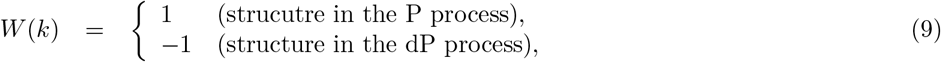

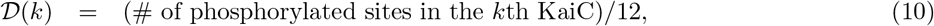

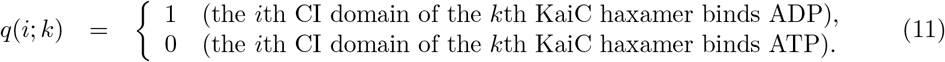

We consider the system to consist of *N* KaiC hexamers. Then, the system state at time *t* is described by a set of vectors, 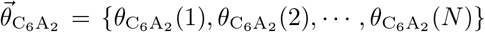, etc. The stochastic evolution of the system state is described by the probability distribution,

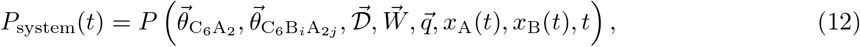

where *x*_A_(*t*) and *x*_B_(*t*) are concentrations of free KaiA dimer and KaiB monomer unbound from KaiC, respectively. The structural change takes place in milliseconds or so in usual protein oligomers, and we assume a similar timescale in the present problem. This timescale is much shorter than the timescales of other reactions; therefore, we treat the variable 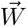 as in quasi-equilibrium. Then, the quasi-equilibrium free energy *G*_quasi_ should be expanded by 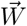 as *G*_quasi_ = *G*_0_ + ∑_*k*_ *W*(*k*)*G*_1_(*k*)+ ∑∑_*k*≠*l*_ *W*(*k*)*W*(*l*)*G*_2_(*k, l*) + ⋯. Here, because *W*(*k*)^2^ = 1, the second-order term of the expansion only consists of the sum with *k* ≠ *l*. In the present model, the interaction between different KaiC hexamers is indirect through the KaiA sequestration; therefore, we can put *G*_2_(*k,l*) = 0. Then, in the expression *G*_quasi_ = *G*_0_ + ∑_*k*_ *W*(*k*)*G*_1_(*k*), *W*(*k*) behaves like the Ising spin under the external field *R*(*k,t*) = –*G*_1_(*k*). Therefore, the average of *W*(*k*) in quasi-equilibrium is 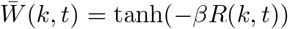. By intoducing the order parameter of structure, 0 ≤ *X*(*k,t*) < 1, as 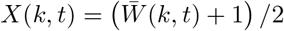, we have

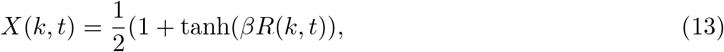

showing the same form as in Eq. 4 in the main text. Here, *β* = 1/(*k*_B_*T*) is inverse temperature, and *R*(*k, t*) is a quasi-equilibrium constraint reflecting the chemical state of the KaiC hexamer at time *t*. The explicit form of *R*(*k,t*) represents how the feedback relations among reactions and structure work in the system and is defined after the other variables in Eq. 12 are transformed to an expression suitable to be handled.

We use the mean-field approximation; Eq. 12 is approximated by factorizing *P* into each degree of freedom as

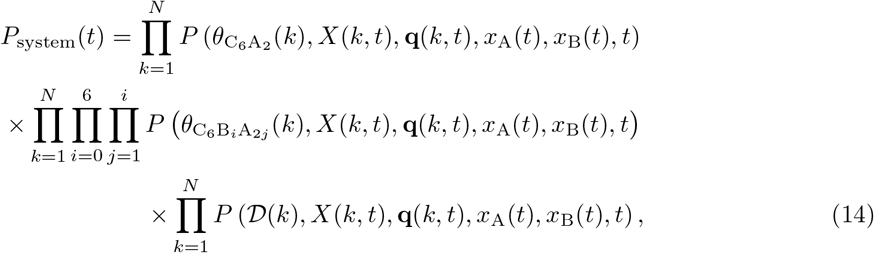

where **q**(*k,t*) = {*q*(1; *k,t*),*q*(2; *k, t*),…, *q*(6; *k,t*)} is a six-dimensional vector representing the nucleotide-binding state of each of six CI domains in the *k*th hexamer. We represent the nonequilibrium consumption of ATP by treating *q*(*i*; *k, t*) as a stochastic variable taking the value either 1 or 0 depending on whether the CI domain binds ADP or ATP (Eq. 11). *X*(*k, t*), *x*_A_(*t*), and *x*_B_(*t*) are calculated by solving the relations explained in the subsections “Reaction-structure feedback coupling” and “Coupling of multiple oscillators” in this Methods section. Thus, by dropping the variables, **q**(*k, t*), *X*(*k, t*), *x*_A_(*t*), and *x*_B_(*t*) from the expression, Eq. 14 is

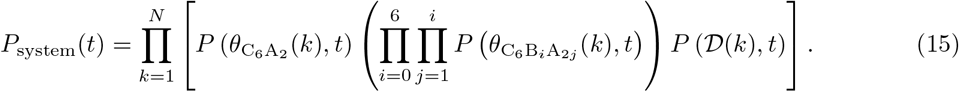

*P*_system_(*t*) should obey the master equation representing reactions in the Kai system. In the mean-field approximation, the master equation is reduced to simpler equations similar to the chemical kinetics equations [74, 75]. Thus, we consider the equation of KaiA binding/unbinding kinetics for *P*(*θ*_c_6_A_2__(*k*),*t*), the equations of KaiB and KaiA binding/unbinding kinetics for *P*(*θ*_C_6_B_*i*_A_2*j*__(*k*),*t*), and the equation of P/dP kinetics for 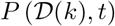.

#### KaiA and KaiB binding/unbinding kinetics

Writing *P*_C_6_A_2__ (*k,t*) = *P*(*θ*_C_6_A_2__(*k*) = 1,*t*), we have the kinetic equation for the KaiA binding and unbinding,

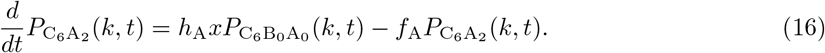

Because the KaiA binding and unbinding reactions are faster than the other reactions in the present system, we can approximate Eq. 16 with the quasi-equilibrium approximation as 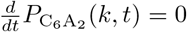. Then, we have

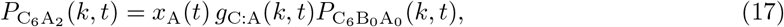

with *g*_C:A_(*k,t*) = *h*_A_(*k,t*)/*f*_A_(*k,t*). As discussed in the main text, *h*_A_ should be an increasing function of *X*(*k,t*) and *f*_A_ is a decreasing function of *X*(*k,t*). We use the form,

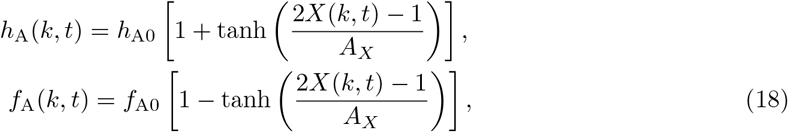

where *h*_A0_ and *f*_A0_ are the rate cosnstants to determine the time scale and *A_X_* > 0 is a constant to determine the sensitvity to the structure.

In a similar way, by writing *P*_C_6_B_*i*_A_2*j*__(*k, t*) = *P*(*θ*_C_6_B_*i*_A_2*j*__(*k*) = 1, *t*), the binding and unbinding of KaiA to and from KaiB should be in quasi-equilibrium, leading to

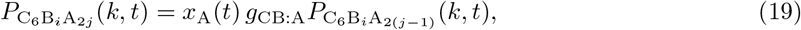

with *g*_CB:A_ = *h*_AB_/*f*_B_, where *h*_AB_ and *f*_AB_ are constants independent of *X*(*k,t*). Then, we can write

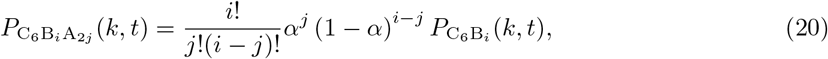

with 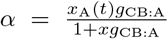, and 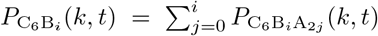 for *i* ≥ 1. Using the variables *P*_C_6_B_*i*__(*k,t*), the kinetic equation for *P*_C_6_B_*i*_A_2*j*__(*k,t*) becomes

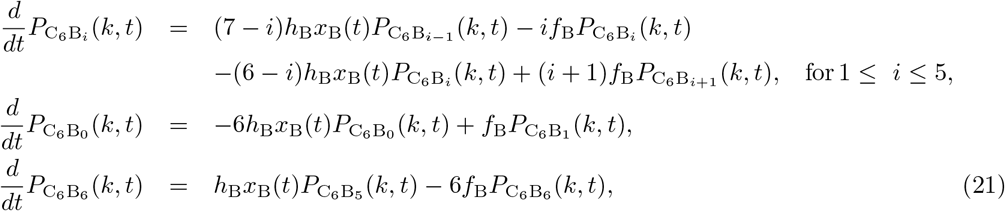

where *P*_C_6_B_0__(*k,t*) = *P*_C_6_A_2__(*k,t*) + *P*_C_6B_0__A_0__(*k,t*), and the rate constants are

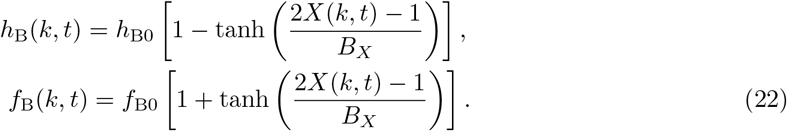

*h*_B0_, and *f*_B0_ are the rate constants defining the time scale and *B_X_* > 0 is a constant determining the sensitivity to the structure. Because the bindable conformation of KaiB appears with the thermal activation with the energy Δ*E*_fs→gs_, we assume *h*_B0_ and *f*_B0_ at temperature *T* are

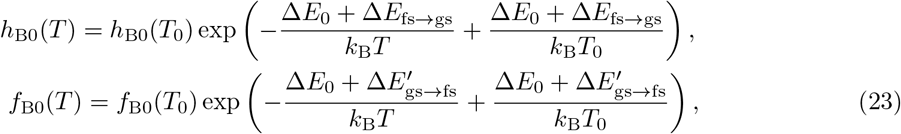

as explained in Table 1 in the main text.

#### P/dP kinetics

For the P/dP reactions, we calculate 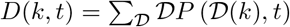. Then, we have

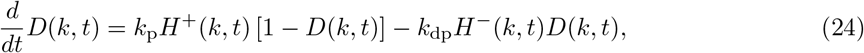

with the rate constants *k*_p_ and *k*_dp_. Here, *H*^+^(*k,t*) = *z*/(1 + *z*) and *H*^-^(*k,t*) = 1/(1 + *z*) are the effects of binding and unbinding of KaiA to and from the CII, respectively, and *z* = *P*_C_6_A_2__(*k, t*)/*P*_0_ with a constant *P*_0_.

#### ATPase reactions

We describe the non-equilibrium ATPase reactions by using a stochastic variable *q*(*i*; *k,t*). When ATP is bound on the ith CI domain of the *k*th KaiC hexamer, we write *q*(*i*; *k, t*) = 0. The ATP hydrolysis is represented by the transition from *q*(*i*; *k,t*) = 0 to *q*(*i*; *k,t*) = 1, which takes place at a random timing with the frequency *f*_hyd_. We approximate that *f*_hyd_ does not depend on *X*, as explained in the main text. The state *q*(*i*; *k,t*) = 1 represents the ADP-bound state. It is not known how the release of inorganic phosphate (P_i_) impacts the KaiC structure. In the present study, for simplicity, we do not distinguish the ADP+P_i_ bound state just after the hydrolysis and the ADP bound state after the P_i_ release. We assume that the lifetime of the ADP bound state 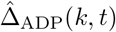 is stochastically fluctuating as

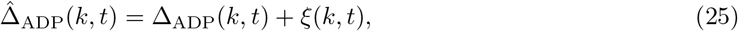

where *ξ*(*k, t*) is a random number satisfying 〈*ξ*(*k, t*)〉 = 0 and 〈*ξ*(*k, t*)(*k*′, *t*′)〉 = *δ_kk′_δ*(*t-t*′)Δ_ADP_(*k, t*). After the ADP release, the ATP rebinds, which turns the nucleotide-binding state from *q*(*i*; *k, t*) = 1 to 0. As explained in the main text, we assume

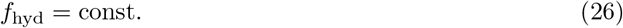

and

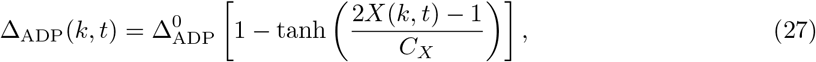

where *C_X_* and 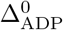 are constants.

The ensemble-averaged rate of the ADP release, 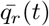, shown in Fig. 2B, was calculated as

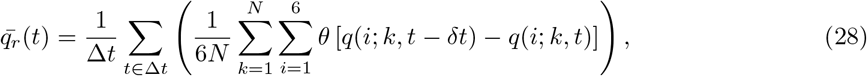

where *θ*(*x*) = 1 for *x* > 0 and *θ*(*x*) = 0 for *x* ≤ 0. *δt* = 10^-3^ h is the width of the simulation time step, and Δ*t* = 0.2 h is the time window for the data sampling.

#### Reaction-structure feedback coupling

*R*(*k,t*) in Eq. 13 represents the major assumptions on the feedback coupling in the present model. We use 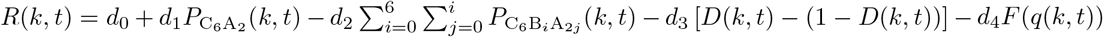 with constants *d*_0_, *d*_1_, *d*_2_, *d*_3_, and *d*_4_, and

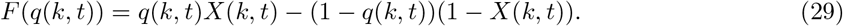

Then, we have

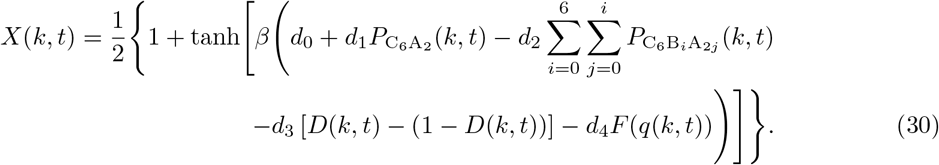

#### Coupling of multiple oscillators through conservation of molecules

The constraints coming from the conservation of the total concentrations of KaiA (Eq. 5 of the main text) and KaiB are simplified with the quasi-equilibrium treatment of KaiA (Eqs. 17 and 19) as

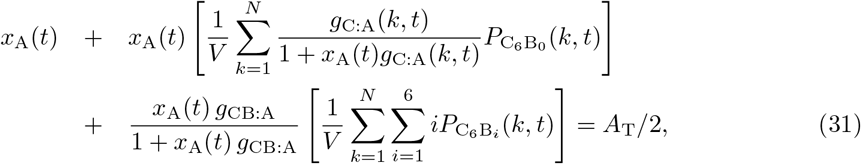

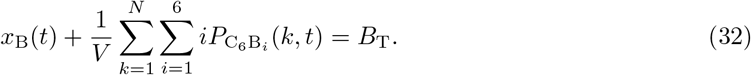

### Simulations

We simulated the system containing *N* = 1000 or 2000 KaiC hexamers by numerically integrating the kinetics with a time step of *δt* = 10^-3^ h. The variables describing the system are *P*_C_6_A_2__(*k,t*), *P*_C_6_B_*i*_A_2*j*__(*k,t*) (0 ≤ *j* ≤ *i* ≤ 6), *q*(*i*; *k,t*) (1 ≤ *i* ≤ 6), *D*(*k,t*), *X* (*k,t*), for *k* = 1 ~ *N*, *x*_A_(*t*) and *x*_B_(*t*). From given values of these variables at time *t*, the values at *t* + *δt* were obtained by (i) stochastically updating *q*(*i*; *k,t*) using a constant *f*_hyd_ or Eqs. 25 and 27, (ii) evaluating the binding and unbinding constants of Eqs. 18 and 22, (iii) updating *P*_C_6_A_2__(*k,t*) and *P*_C_6_B_*i*_A_2*j*__(*k,t*) with Eqs. 17 and 21, (iv) updating *x*_A_ and *x*_B_ by solving Eqs. 31 and 32, (v) updating *D*(*k, t*) with Eq. 24, and (vi) updating *X*(*k, t*) using Eq. 30. The period and amplitude were calculated from the trajectories, each having the length 3276.8 h obtained after the initial warming-up trajectories of 100 h length.

### Parameters

The KaiC concentration is *C*_T_ = 3.3*μ*M on a monomer basis for *N* = 1000 and *V* = 3 × 10^-15^*l*; this concentration is close to 3.5*μ*M, often used in experiments. We assume the ratio *A*_T_ : *B*_T_ : *C*_T_ = 1 : 3 : 3 as in many experiments [67, 23, 54]. The oscillations were robust against small parameter changes; therefore, we did not calibrate the parameters but determined them from the order of magnitude argument. We chose *h*_B0_*B*_T_ and *f*_B0_ to satisfy *h*_B0_*B*_T_ ≈ *f*_B0_ ≈ 1 h^-1^. We used *h*_B0_ = 5 × 10^-5^ h^-1^ and *f*_B0_ = 2 h^-1^ in units of *V* = 1, corresponding to the dissociation constant of 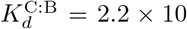 *μ*M in *V* = 3 × 10^-15^*l*. We used *h*_A0_/*f*_A0_ = 5 × 10^-4^ in units of *V* = 1, corresponding to 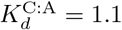 *μ*M in *V* = 3 × 10^-15^*l*, which agrees with the experimentally observed values of 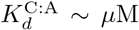 [76]. The dissociation constant of KaiA and KaB was not yet observed experimentally. Here, we assumed a smaller value to ensure the sequestration effect; *g*_CB:A_ = 6 × 10^-3^ in units of *V* = 1, corresponding to 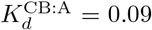 *μ*M in *V* = 3 × 10^-15^*l*. See Tables 2 and 3 for values of the other parameters. We chose *d*_0_, *d*_1_, *d*_2_, *d*_3_. and *d*_4_ as of the order of *k*_B_*T*_0_, and *A*_X_, *B*_X_, and *C*_X_ as of the order of one.

**Table 3:**
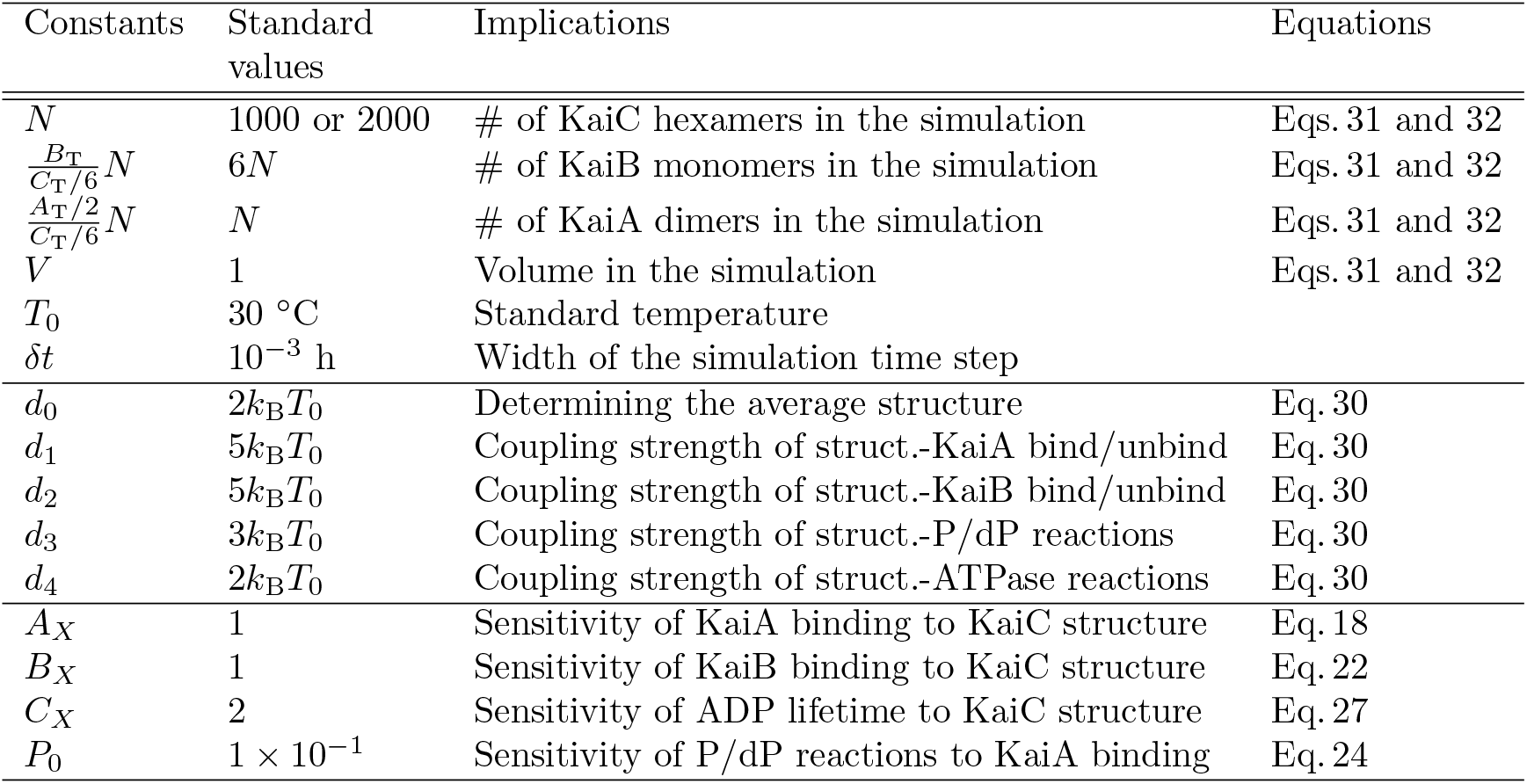
Other parameters.

## Acknowledgments

This work was supported by JSPS-KAKENHI Grants JP20H05530, 21H00248, and 22H00406.

